# The atypical adhesion GPCR ADGRA1 controls hippocampal inhibitory circuit function

**DOI:** 10.1101/2025.07.30.667713

**Authors:** Baris Tosun, Elizabeth Orput, Duy Lan Huong Bui, Richard C. Sando

## Abstract

Neural circuits contain a diverse array of inhibitory interneurons that control information processing. The cell surface receptors and signaling pathways that modulate cell type specific inhibitory synaptic function are unclear. Here, we identify the atypical adhesion GPCR ADGRA1 as essential for hippocampal PV and SST inhibitory synaptic function. ADGRA1 is selectively enriched in hippocampal PV and SST interneurons and localizes to a subset of synapses. ADGRA1 deletion in PV and SST interneurons impairs inhibitory synaptic inputs onto Dentate Gyrus granule cells and generates deficits in learning and memory. ADGRA1 engages several downstream G proteins, notably Gα13, a pathway important for the establishment of hippocampal PV interneuron synaptic networks. These results identify an orphan receptor pathway selective for specific inhibitory synapse subtypes and expand our understanding of the signaling mechanisms that establish hippocampal inhibitory circuits.

## Introduction

Mammalian neural circuitry contains an array of physiologically and morphologically diverse inhibitory interneurons that are critical modulators of information processing^1^. While representing only ∼10-15% of the total hippocampal neuron population, interneurons shape circuit function via networks of synapse specific connectivity^2^. Their cell type and synapse specific functions are essential for learning and memory. Moreover, experience and neuronal activity shape the plasticity of interneuronal synapses in a cell type specific manner^3^. Interneuron dysfunctions are highly associated with neurological disorders including autism spectrum disorder (ASD), Rett syndrome and epileptic encephalopathies^4^. Despite the importance of diverse inhibitory synaptic connections, the bi-directional synaptic signaling pathways that control the establishment of these cell type specific GABAergic synapses remain unclear.

Increasing evidence supports the importance of adhesion G protein-coupled receptors (aGPCRs) in synapse formation and neural circuit assembly^5–9^. aGPCRs are a large GPCR class that display both extracellular adhesion and intracellular signaling functions^10,11^. A signature feature of the aGPCR class is a membrane proximal GAIN (GPCR autoproteolysis inducing) domain^12^. The GAIN domain has been a focus of aGPCR signal activation mechanisms^13,14^. The GAIN domain of many aGPCRs exhibits an autoproteolysis site which can generate a membrane-proximal self-tethered agonist (TA), also known as a *Stachel* peptide^15^. The TA can induce signaling in the C-terminal fragment (CTF) GPCR via either removal of the N-terminal fragment (NTF) and exposure, or through a more tunable mechanism involving the NTF^16^.

Of the 33 human aGPCRs, all exhibit a GAIN domain with one exception, ADGRA1 (GPR123)^10^. ADGRA1 lacks N-terminal extracellular adhesion domains together with the GAIN, suggesting unique functions and signaling mechanisms compared to other aGPCRs. ADGRA1 exhibits a 7-transmembrane (7-TM) GPCR followed by a relatively large cytoplasmic tail (268 amino acids in mice) with a C-terminal PDZ binding motif but lacks extensive extracellular regions characteristic of other aGPCRs. Despite these unique features, ADGRA1 is evolutionarily conserved and exhibits 37% sequence identity with ADGRA2 (GPR124) and 44% with ADGRA3 (GPR125) in the 7-TM region and C-terminal tail. Unlike ADGRA1, ADGRA2 and ADGRA3 contain extensive extracellular adhesion modules composed of leucine rich repeats and immunoglobulin domains^11^.

Despite these interesting properties, the biological functions of ADGRA1 remain unclear. Elevated *Adgra1* expression has been identified in human bladder and breast cancer studies^17,18^, and has been shown to regulate maintenance and acquisition of pluripotency in hiPSCs^19^. *Adgra1* is highly expressed in the mammalian CNS^20^, and proteomic analysis found it enriched in postsynaptic fractions, supporting synaptic localization^21^. Functional studies using constitutive Adgra1 KO mice found reduced body weight and deficiencies in metabolism and thermogenesis, likely due to dysfunctions in the hypothalamus^22^. Further analysis of constitutive KO mice determined altered anxiety-like behaviors in mutant male mice^23^. However, the cell type-specific and putative synaptic role of ADGRA1 in mammalian circuits remains unknown.

Our studies identify ADGRA1 as highly enriched in hippocampal PV and SST interneurons. ADGRA1 deletion selectively from these GABAergic cell types impairs inhibitory synaptic function and learning and memory. ADGRA1 activates several G proteins, notably Gα13, an important signaling pathway for inhibitory synaptic function, and localizes with Gα13 at synapses. Collectively, these studies identify a synaptic orphan receptor that controls inhibitory synaptic function in a cell type specific manner.

## Results

### ADGRA1 is enriched in hippocampal PV- and SST-positive interneurons and localizes to synapses

To examine aGPCR expression in hippocampal inhibitory neurons, we crossed Cre-inducible Ribotag (HA-Rpl22) mice to the vGAT-Cre driver line (**Fig. 1A-C**). This approach enables immunoprecipitation of ribosomal-bound mRNAs in a cell type-specific manner^24^. We performed HA immunoprecipitations from postnatal day 30 (P30) hippocampal tissue and analyzed the enrichment of aGPCRs and control transcripts relative to input mRNA samples using RT-qPCR (**Fig. 1A-C**). Interestingly, of the aGPCRs tested, *Adgra1* was enriched in hippocampal interneurons (**Fig. 1B**). Controls transcripts confirmed the enrichment of inhibitory markers in immunoprecipitated samples (*PV*, *SST*, and *Slc32a1/vGAT*) and de-enrichment of glial (*Gfap*) and excitatory neuronal (*Slc17a7/vGlut1*) markers (**Fig. 1C**). Next, we conducted RNA *in situ* hybridizations to examine the spatial expression pattern of *Adgra1* relative to other cell type molecular markers (**Fig. 1D-G**, **Fig. 2A& B, Fig. S1, S2**). *Adgra1* was highly expressed in sparse sub-populations of cells throughout the P30 hippocampus (**Fig. 1D & E**). We analyzed the developmental time course of *Adgra1* expression throughout the hippocampus and found that it was expressed at negligible levels shortly after birth and increased during postnatal development (**Fig. S1A & B**). We next conducted double RNA *in situ*/immunohistochemistry experiments to determine if the sparse population of *Adgra1* expressing cells are neurons or glia (**Fig. S1C-F**). *Adgra1* expression overlapped with the neuronal marker NeuN but not GFAP, supporting high expression in a sub-population of hippocampal neurons (**Fig. 1C-F**). These results support that *Adgra1* is an aGPCR enriched in hippocampal interneurons.

**Figure 1:**
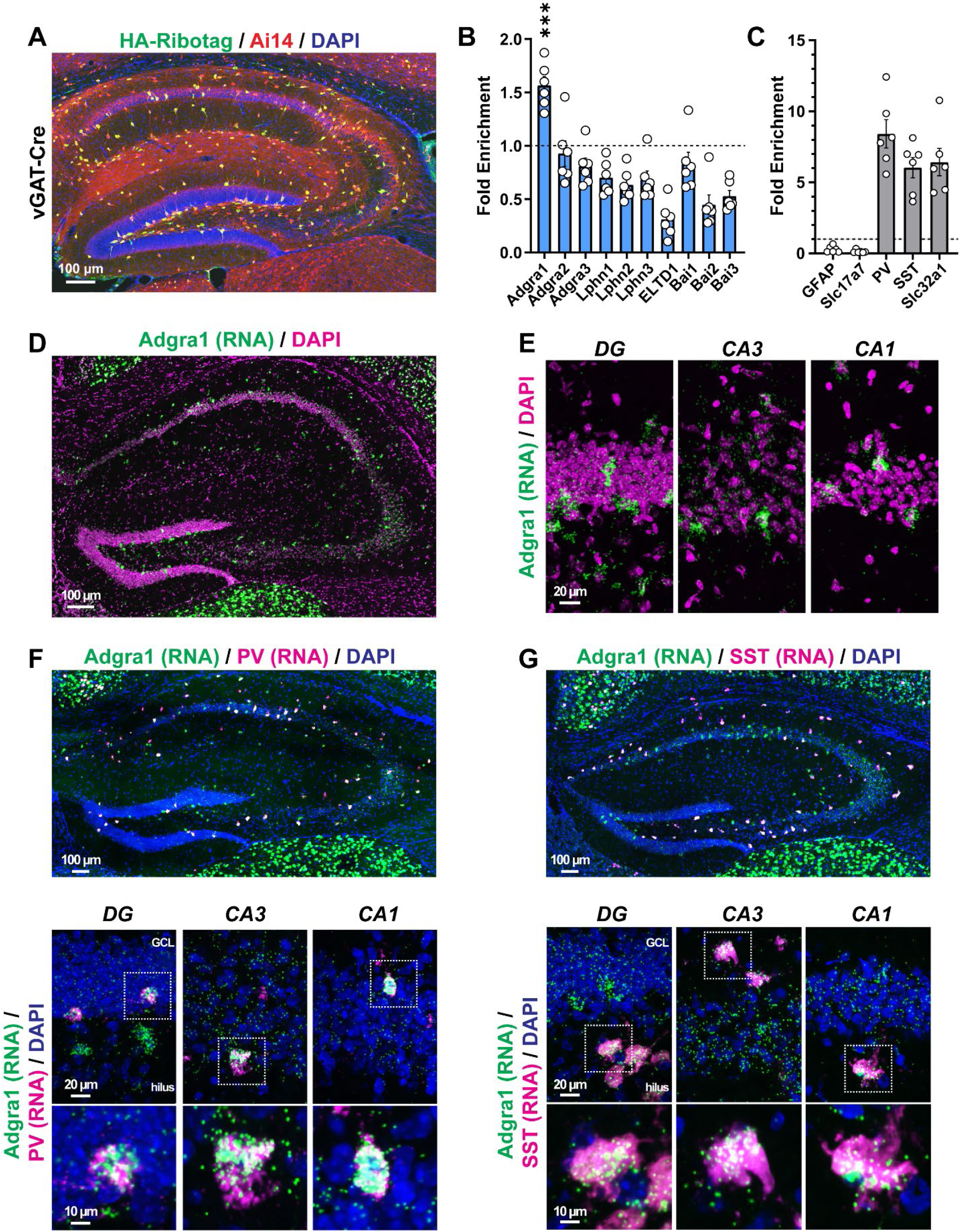
The atypical orphan aGPCR Adgra1 is enriched in hippocampal interneurons. **A,** representative sections from mice harboring the Cre-inducible RiboTag (HA-Rpl22) allele, vGAT-Cre, and the Ai14 reporter immunolabeled for HA together with DAPI. **B,** RiboTag experiments from ribosomal bound mRNA from postnatal day (P30) hippocampus for indicated aGPCRs. The relative enrichment of each target in the HA RiboTag immunoprecipitation was compared to the input sample from the same tissue. **C,** qPCR controls verifying enrichment of interneuronal transcripts from vGAT-Cre RiboTag experiments. Probes for interneurons (*Pvalb*, *SST*, *Slc32a1/vGAT*) were compared to probes for glial (*Gfap*) or excitatory neuron (*Slc17a7/vGlut1*) transcripts. **D & E,** *Adgra1* is expressed in sparse subpopulations of cells in the postnatal hippocampus. **D,** representative RNA *in situ* of the P30 hippocampus labeled for *Adgra1* together with DAPI. **E,** representative high-magnification images of RNA *in situs* for *Adgra1* in the dentate gyrus (DG), CA3, and CA1. **F & G,** *Adgra1* is enriched in hippocampal PV and SST-positive interneurons. **F,** RNA *in situ* hybridizations for *Adgra1* together with *PV*. **G,** similar to **F**, except for *in situ* hybridizations for *Adgra1* and *SST*. Numerical data are means ± SEM from 6 independent biological replicates (mice). See Figure S1 and S2 for additional analysis of *Adgra1* expression in the brain.

**Figure 2:**
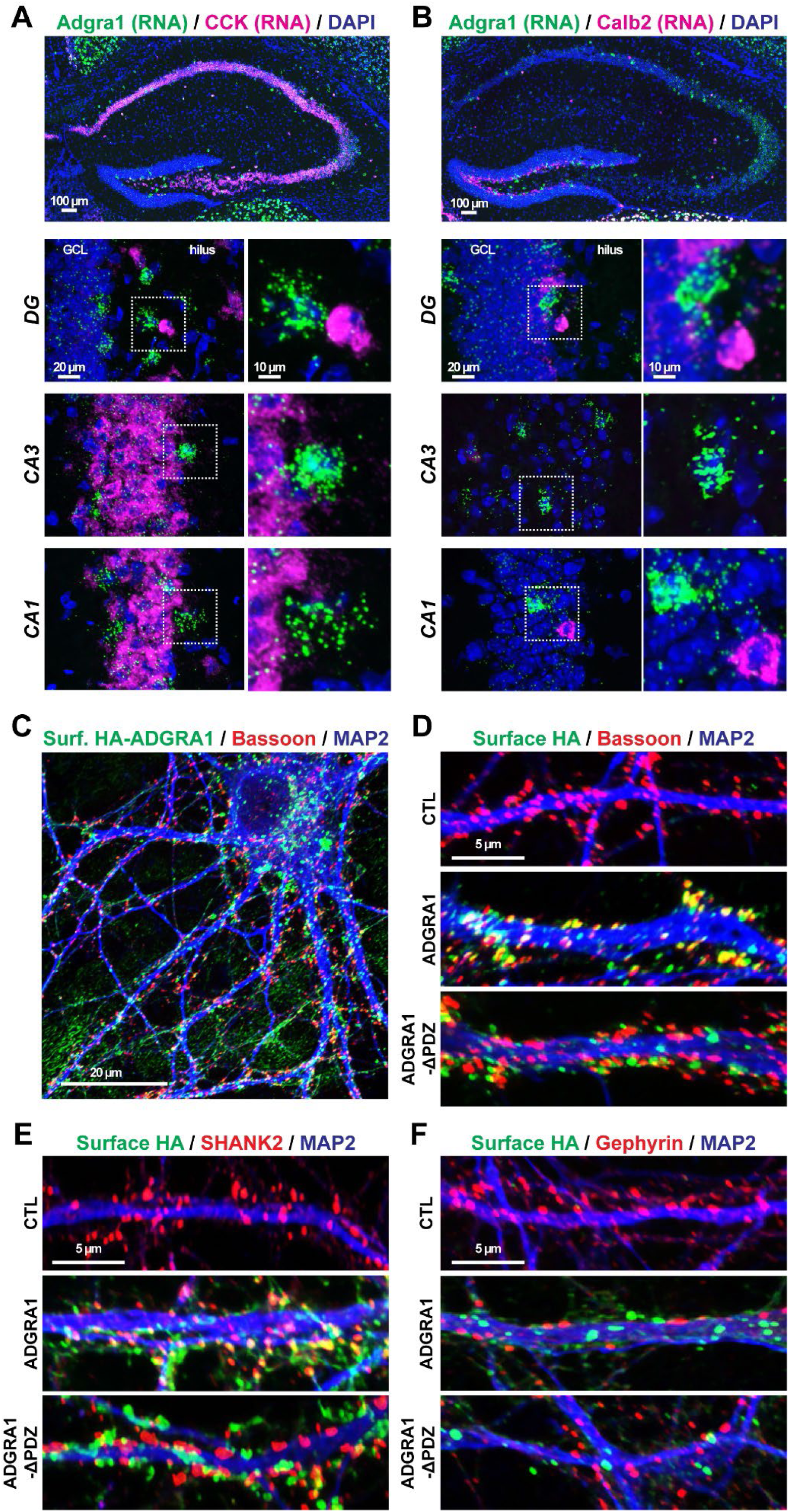
*Adgra1* is selectively expressed in PV- and SST-positive hippocampal interneurons and is a synaptic GPCR. **A & B,** *Adgra1* is absent from hippocampal CCK- and Calb2-positive interneurons. **A,** representative RNA *in situ* for *Adgra1* together with *CCK* in the P30 hippocampus. **B,** similar to **A**, except for RNA *in situs* probing for *Adgra1* and *Calb2*. **C-F,** immunocytochemistry for HA-tagged ADGRA1 in primary hippocampal neurons. **C,** representative primary hippocampal neurons transduced with lentivirus expressing HA-ADGRA1 and immunolabeled for surface HA followed by presynaptic Bassoon and the somatodendritic marker MAP2. **D,** example dendrite immunolabeled for surface HA-ADGRA1 followed by Bassoon and MAP2. **E,** immunocytochemistry for surface HA-ADGRA1 and intracellular SHANK2 and MAP2. **F,** immunocytochemistry for surface HA-ADGRA1 and intracellular Gephyrin and MAP2. See Figure S1 and S2 for additional analysis of *Adgra1* expression in the brain and Figure S3 for quantification of ADGRA1/synaptic marker colocalization.

To determine the subtype of interneuron, we performed RNA *in situ* experiments in the P30 hippocampus for *Adgra1* together with interneuronal markers including *PV*, *SST*, *CCK*, and *Calb2* (**Fig. 1F & G, Fig. 2A & B, Fig. S2**). *Adgra1* expression highly co-localized with *PV* and *SST* (**Fig. 1F & G**). *Adgra1* was excluded from *CCK* and *Calb2* expressing cells, suggesting that *Adgra1* is selective for PV and SST inhibitory subtypes (**Fig. 2A & B**). *Adgra1* was also expressed in distinct patterns in other brain regions including cortical layer 6 and the thalamus and therefore may also have important functions in these areas (**Fig. S2**). These results show that *Adgra1* is an orphan aGPCR selectively expressed in hippocampal PV and SST interneurons.

We next assessed the subcellular localization of ADGRA1 in hippocampal neurons. Given the absence of reliable antibodies for ADGRA1, we expressed HA-tagged ADGRA1 via lentivirus in primary hippocampal cultures (**Fig. 2C-F, Fig. S3**). We compared the cell surface localization of WT ADGRA1 to a PDZ binding motif truncation (ADGRA1-ΔPDZ) to determine if this sequence is important for localization. Surface ADGRA1 formed puncta that partially co-localized with synapses (**Fig. 2C-F, Fig. S3**). ADGRA1-ΔPDZ localized to synapses comparable to WT, suggesting that other sequence features are responsible for synaptic localization. Collectively, these results show that ADGRA1 is a synaptic GPCR selectively enriched in hippocampal PV and SST interneurons.

### ADGRA1 is dispensable for synaptic function in excitatory DG GCs

We subsequently sought to elucidate the function of ADGRA1 in PV and SST hippocampal interneurons. We generated Adgra1 floxed mice (Adgra1 cKO) and validated this allele by generating primary hippocampal cultures and tranducing them with lentiviral Cre or inactive ΔCre as a control (**Fig. S4**). This approach enables high efficiency of Cre delivery to obtain a robust KO across the cell population. Adgra1 cKO cultures infected with Cre lacked floxed exon 6 relative to ΔCre conditions, supporting effective deletion with this allele (**Fig. S4**). Transcripts for other aGPCRs were unaltered (**Fig. S4**). Next, we crossed Adgra1 cKO models to several Cre drivers to obtain cell type-specific deletion. First, since we observed low levels of *Adgra1* expression throughout the hippocampal granule cell and pyramidal cell layers (**Fig. 1**), we crossed Adgra1 cKO mice to the EMX1-Cre driver to delete ADGRA1 from forebrain excitatory neurons and glia (**Fig. 3, Fig. S5, S6**)^25^. In parallel, we crossed Adgra1 cKO models to the PV-Cre or SST-Cre driver lines to delete ADGRA1 specifically in PV or SST interneurons (**Fig. 4-6, Fig. S7-S10**). We focused on the granule cell (GC) circuitry of the dentate gyrus (DG) given the importance of PV and SST inhibitory synaptic function in this area as well as its well characterized patterns of synaptic connectivity^26,27^.

**Figure 3:**
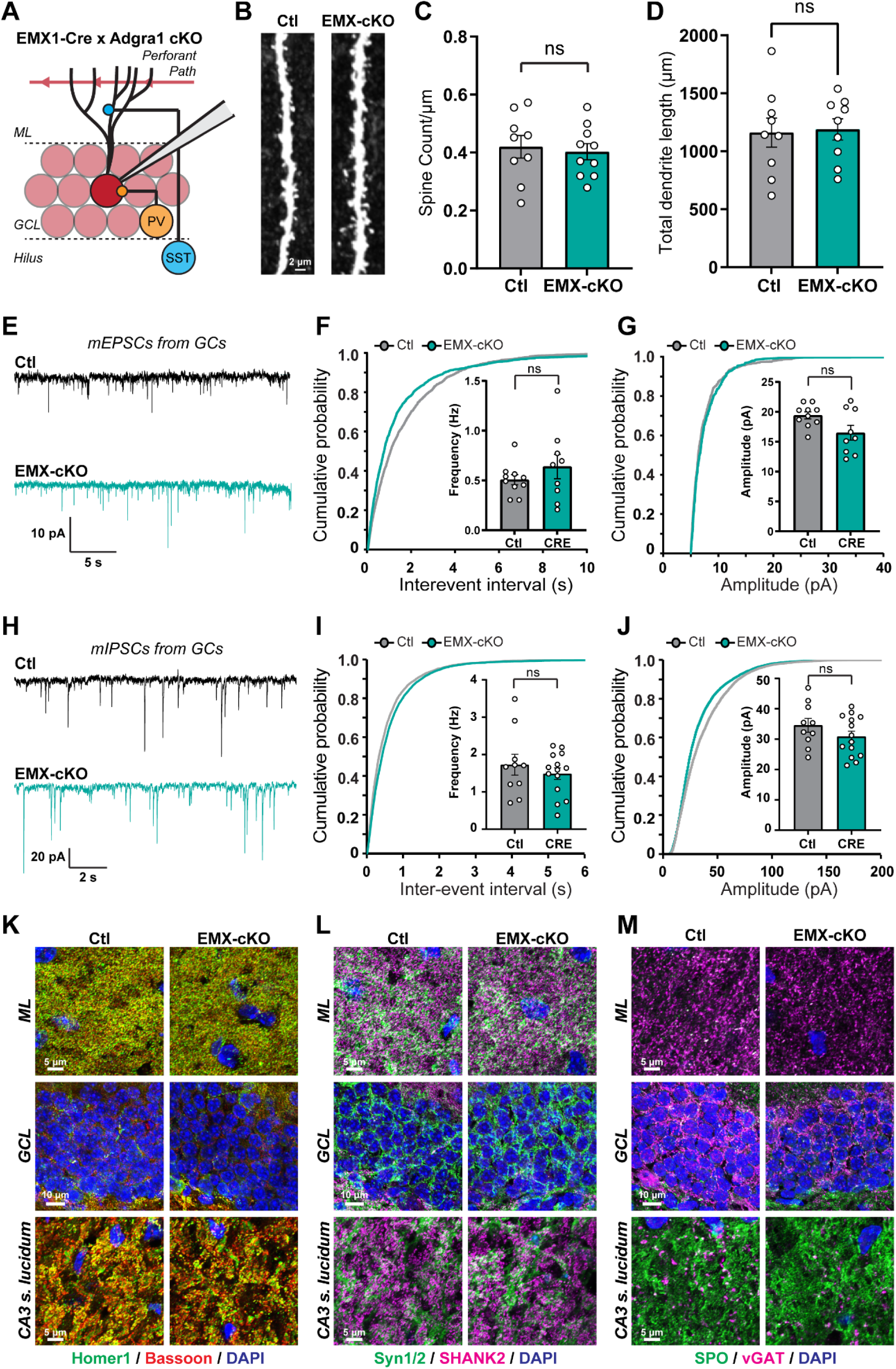
Neuronal morphology, synapse density, and synaptic transmission are preserved in dentate gyrus (DG) granule cells (GCs) lacking ADGRA1. **A,** experimental diagram focused on the hippocampal DG circuit. Adgra1 cKO mice were crossed to the EMX1-Cre driver line to delete ADGRA1 from excitatory neurons. **B-D,** morphological analyses of DG GCs. GCs were filled with biocytin during electrophysiological recordings and dendritic spine density and dendritic arborization subsequently analyzed. **B,** representative spine images from Ctl or EMX-cKO GCs. **C,** quantification of dendritic spine densities in GCs. **D,** quantification of average total dendrite length from DG GCs. **E-G,** analysis of mEPSCs from Ctl or EMX-cKO GCs. **E,** representative mEPSC traces. **F,** cumulative probability plot of inter-event intervals and summary graph [inset] of the mean mEPSC frequency. **G,** cumulative probability plot and summary graph [inset] of mEPSC amplitude measurements. **H-J,** analysis of mIPSCs from Ctl or EMX-cKO GCs. **H,** representative mIPSC traces. **I,** cumulative probability plot of inter-event intervals and summary graph [inset] of the mean mIPSC frequency. **J,** cumulative probability plot and summary graph [inset] of mIPSC amplitude measurements. **K-M,** immunohistochemical analysis of synapses in the DG. **K,** immunolabeling for presynaptic Bassoon together with postsynaptic excitatory Homer1 in the hippocampal DG molecular layer (ML), granule cell layer (GCL), or CA3 stratum lucidum. **L,** similar to **K**, except for immunolabeling for presynaptic Syn1/2 and postsynaptic excitatory SHANK2. **M,** similar to **K**, except for presynaptic inhibitory vGAT or synaptoporin (SPO), a marker of DG large mossy fiber terminals (LMTs). Numerical data are means ± SEM or cumulative histograms. See Figure S4 for characterization of the Adgra1 cKO mouse allele, Figure S5 for additional electrophysiological and morphological parameters, and Figure S6 for quantification of immunohistochemistry. Control (Ctl) mice were Adgra1 cKO and EMX-cKO were EMX1-Cre/Adgra1 cKO littermates. Statistical significance was determined via two-tailed t-test or Kolmogorov-Smirnov test (cumulative histograms).

**Figure 4:**
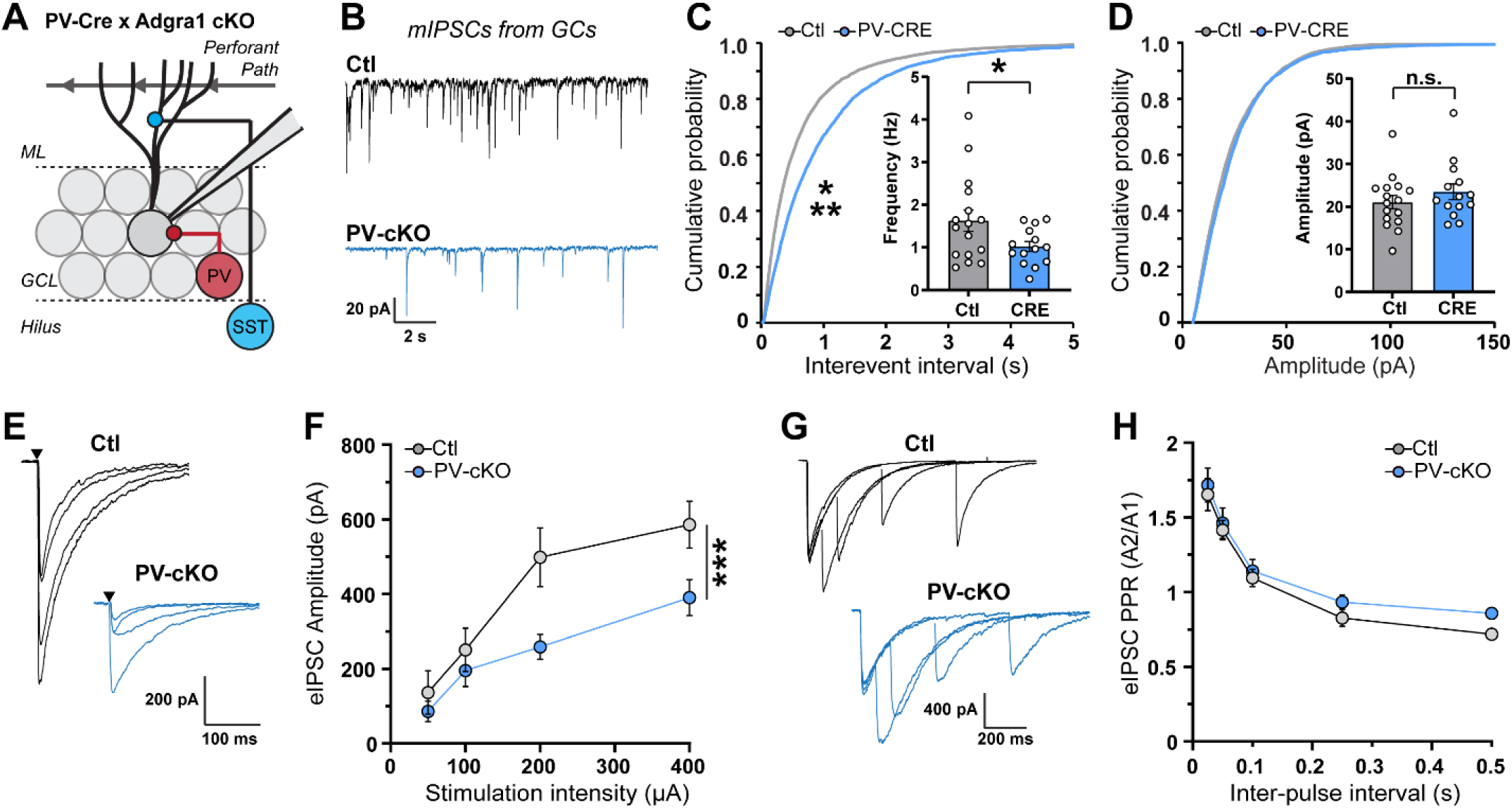
ADGRA1 is essential for inhibitory synaptic function in hippocampal PV interneurons. **A,** diagram of experimental approach to determine ADGRA1 function in PV inhibitory DG circuits. **B-D,** analysis of spontaneous mIPSCs from Ctl or PV-cKO GCs. **B,** representative mIPSC traces from GCs. **C,** cumulative probability plot of inter-event intervals and summary graph [inset] of the mean mIPSC frequency. **D,** cumulative probability plot and summary graph [inset] of mIPSC amplitude measurements. **E-H,** evaluation of eIPSCs in Ctl or PV-cKO GCs. **E,** representative eIPSC traces. **F,** eIPSC amplitude in response to increasing extracellular stimulation. **G,** representative eIPSC paired-pulse ratio (PPR) traces. **H,** quantification of PPR from GCs in Ctl or PV-cKO mice. Numerical data are means ± SEM or cumulative histograms. See Figure S7 for characterization of PV-Cre and SST-Cre lines and Figure S8 for additional electrophysiological parameters. Control (Ctl) mice were Adgra1 cKO and PV-cKO were PV-Cre/Adgra1 cKO littermates. Statistical significance was determined via two-tailed t-test or Kolmogorov-Smirnov test (*, p<0.05; ***, p<0.001).

**Figure 5:**
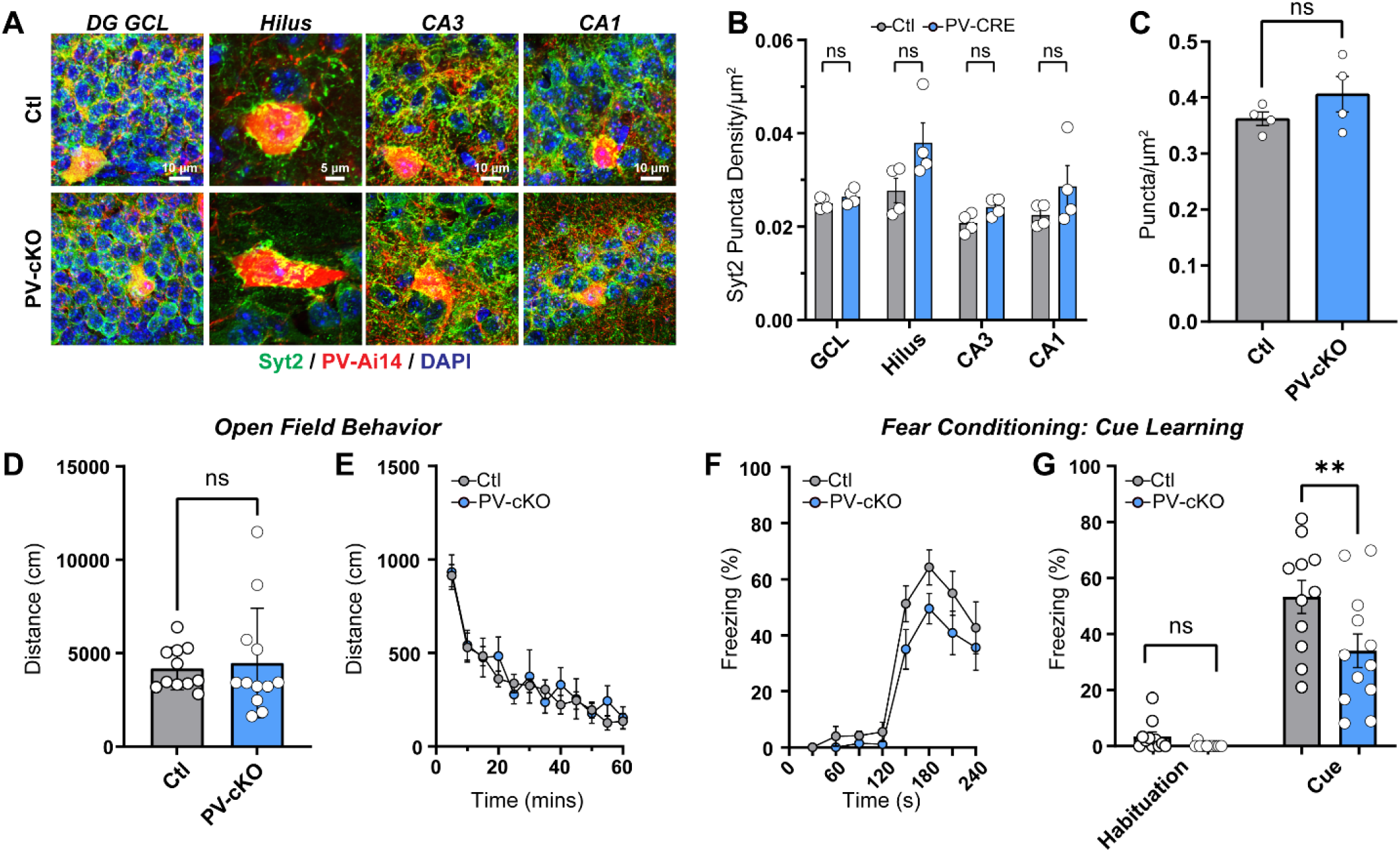
ADGRA1 deletion in PV interneurons impairs learning and memory. **A,** representative images of Syt2-labeled presynaptic PV terminals in the indicated hippocampal sub-regions. **B,** quantification of Syt2 puncta density in the indicated hippocampal sub-regions. **C,** analysis of Syt2-positive puncta density on PV-Ai14 labeled soma. **D & E,** assessment of open field behavior in Ctl or PV-cKO mice. **D,** average distance travelled over a 60-minute open field trial. **E,** distance travelled over time during the 60-minute open field trial. **F & G,** cued learning from Pavlovian fear conditioning studies. **F,** quantification of percent time spent freezing before or after presentation of the cue stimulus following fear conditioning. **G,** percent time spent freezing during the habituation period or presentation of the cue. Numerical data are means ± SEM. See Figure S9 for additional behavioral characterization of PV-cKO mice. Control (Ctl) mice were Adgra1 cKO and PV-cKO were PV-Cre/Adgra1 cKO littermates. Statistical significance was determined via two-tailed t-test, one-way ANOVA with *post hoc* Tukey test or two-way ANOVA (**, p<0.01).

**Figure 6:**
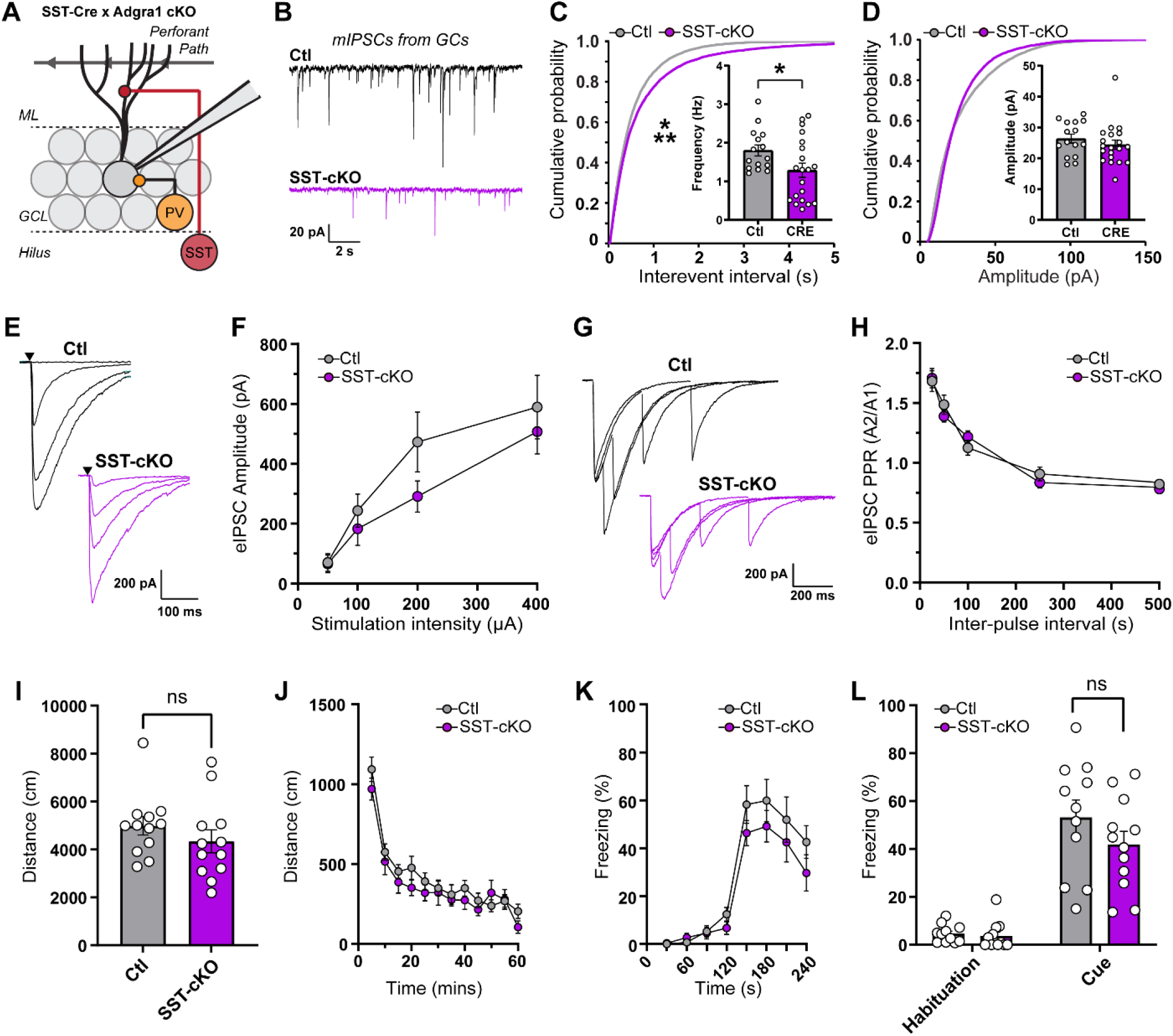
Adgra1 deletion in SST interneurons reduces inhibitory input onto DG GCs. **A,** diagram of experimental approach to determine ADGRA1 function in SST inhibitory DG circuits. **B-D,** analysis of spontaneous mIPSCs from Ctl or SST-cKO GCs. **B,** representative mIPSC traces from GCs. **C,** cumulative probability plot of inter-event intervals and summary graph [inset] of the mean mIPSC frequency. **D,** cumulative probability plot and summary graph [inset] of mIPSC amplitude measurements. **E-H,** eIPSC measurements in Ctl or SST-cKO GCs. **E,** representative eIPSC traces. **F,** eIPSC amplitude in response to increasing extracellular stimulation. **G,** representative eIPSC paired-pulse ratio (PPR) traces. **H,** quantification of PPR from GCs in Ctl or SST-cKO mice. **I & J,** assessment of open field behavior in Ctl or SST-cKO mice. **I,** average distance travelled over a 60-minute open field trial. **J,** distance travelled over time during the 60-minute open field trial. **K & L,** cued learning from Pavlovian fear conditioning. **K,** quantification of percent time spent freezing before or after presentation of the cue stimulus following fear conditioning. **L,** percent time spent freezing during the habituation period or presentation of the cue. Numerical data are means ± SEM or cumulative histograms. See Figure S10 for additional electrophysiological parameters and Figure S11 for additional behavior studies. Control (Ctl) mice were Adgra1 cKO and SST-cKO were SST-Cre/Adgra1 cKO littermates. Statistical significance was determined via two-tailed t-test or Kolmogorov-Smirnov test (*, p<0.05; ***, p<0.001).

We performed whole-cell patch-clamp electrophysiology and obtained recordings from GCs in EMX-cKO or Ctl acute brain slices. We first measured miniature excitatory and inhibitory postsynaptic currents (mEPSCs and mIPSCs). We filled cells during recordings with biocytin to enable subsequent 3D reconstruction and morphological analysis of their dendritic arborizations and dendritic spines (**Fig. 3A-J**). GCs lacking ADGRA1 exhibited no detectable alterations in dendritic spine density and dendrite length and complexity (**Fig. 3B-D**). Moreover, EMX-cKO GCs displayed no significant amplitude and frequency changes in spontaneous synaptic transmission (**Fig. 3E-J, Fig. S5**). We subsequently analyzed synaptic density using immunohistochemistry for pre- and postsynaptic markers (**Fig. 3K-M, Fig. S6**). We detected no significant differences in the density or intensity of excitatory or inhibitory synaptic markers as well as Synaptoporin (SPO), a marker for large mossy fiber terminals (LMTs) formed by GCs (**Fig. 3K-M, Fig. S6**). Thus, ADGRA1 has a minor synaptic role within excitatory GCs of the DG.

### ADGRA1 is essential for hippocampal PV inhibitory synaptic function and learning

We next analyzed PV-Cre/Adgra1 cKO (PV-cKO) models via imaging, electrophysiology, and behavior (**Fig. 4 & 5, Fig. S7-9**). While the PV-Cre and SST-Cre drivers are well characterized and routinely used, we first confirmed their specificity (**Fig. S7**). Interestingly, GCs from PV-cKO acute slices displayed significantly decreased mIPSC frequency but not amplitude (**Fig. 4B-D, Fig. S8A**). To discern the effect on inhibitory synaptic strength compared to presynaptic release probability, we next analyzed evoked IPSCs (eIPSCs) and eIPSC paired-pulse ratios (PPR) (**Fig. 4E-H, Fig. S8**). PV-cKO GCs exhibited significantly lower eIPSC amplitudes, particularly at increasing stimulation strengths (**Fig. 4E & F**). However, PV-cKO GCs displayed no changes in PPR or coefficient of variation in eIPSCs, supporting that presynaptic release probability is preserved (**Fig. 4G & H, Fig. S8C**). These results show that ADGRA1 has a critical function in controlling the presynaptic inhibitory strength of PV interneurons onto GCs.

Importantly, despite this decrease in inhibitory strength, the overall density of PV terminals labeled with Synaptotagmin-2 (Syt2) was unaltered throughout the hippocampus, suggesting a functional but not developmental role of ADGRA1 in presynaptic PV interneurons (**Fig. 5A-C**). We then assessed the consequences of ADGRA1 deletion in PV interneurons on hippocampal-dependent learning and memory (**Fig. 5D-G, Fig. S9**). Open field behavior was unaltered in PV-cKO models (**Fig. 5D & E, Fig. S9A & B**). However, ADGRA1 deletion in PV interneurons produced an impairment of Pavlovian fear conditioning (**Fig. 5F & G, Fig. S9C & D**). This deficit was selective for cued learning while contextual learning was unaltered (**Fig. S9C & D**). Given alterations in hippocampal excitatory/inhibitory balance can generate seizures, we analyzed seizure susceptibility using PTZ (pentylenetetrazole)-mediated seizure induction paradigms but found no significant changes (**Fig. S9E-I**), supporting a role of ADGRA1 function in learning and memory.

Given ADGRA1 is also enriched in SST interneurons, we assessed ADGRA1 synaptic function at the SST-GC circuit by generating SST-Cre/Adgra1 cKO mice (**Fig. 6A**). Spontaneous mIPSC frequency and eIPSC amplitude were reduced in GCs, while release probability measured via PPR was unaltered (**Fig. 6B-H, Fig. S10A-C**). Open field behavior, fear conditioning, and seizure susceptibility was also mostly intact in SST-Cre/Adgra1 cKO mice (**Fig. 6I-L, Fig. S11**). Collectively, these results support that ADGRA1 controls cell type-specific inhibitory synaptic strength onto DG GCs.

### ADGRA1 activates several G proteins including Gα13

Of the 33 aGPCRs, ADGRA1 is the only member that lacks an extensive extracellular region harboring adhesion domains and the GAIN domain (**Fig. 7A**). We assessed which G proteins ADGRA1 is capable of activating using the complete panel of TRUPATH BRET2 biosensors (**Fig. 7B**)^28^. Given the agonist for ADGRA1 is unknown, we examined the ability of overexpressed ADGRA1 to activate the complete panel of 14 TRUPATH Gαβγ sensors relative to empty vector transfected controls. Full-length ADGRA1 activated several G proteins, most notably Gα13 (**Fig. 7B**). While the short extracellular sequence of ADGRA1 has no homology to the aGPCR TA, we determined if this extracellular sequence is required for G protein activation by replacing it with a short glycine linker (ΔN-ADGRA1) (**Fig. 7C**). We first determined the relative expression level of overexpressed full-length ADGRA1 compared to ΔN-ADGRA1 using immunocytochemistry and immunoblotting (**Fig. S12**). The ΔN-ADGRA1 mutant was expressed at comparable levels to full-length ADGRA1 (**Fig. S12**). The G protein coupling profile of ΔN-ADGRA1 was similar to full-length ADGRA1, suggesting that this extracellular sequence is not involved in basal G protein activation (**Fig. 7C**). We subsequently validated ADGRA1 G protein coupling by performing full-length ADGRA1 plasmid dose-response experiments to measure the relationship between ADGRA1 plasmid copy number and BRET2 response (**Fig. 7D-G**). GαoB, which displayed no significant change in BRET2 signal (**Fig. 7B**), had no dose-dependent increase in ΔBRET2 (**Fig. 7D**). However, Gα11, Gα15, and Gα13 exhibited a plasmid copy number-dependent change in BRET2, supporting the specificity of these measurements. Our previous studies found that Gα13 partially co-localizes with synaptic markers and has important roles in PV synaptic function^29^. We next examined the co-localization of ADGRA1 with endogenous Gα13 (**Fig. 7H, Fig. S12E**). HA-ADGRA1 puncta partially co-localized with Gα13 puncta in primary hippocampal neurons (**Fig. 7H & I, Fig. S12E)**. Similar to co-localization studies with synaptic markers (**Fig. 2C-F, Fig. S3**), deletion of the PDZ-binding motif preserved this co-localization. Collectively, these experiments show that ADGRA1 activates several G proteins including Gα13 and co-localizes with Gα13 in hippocampal neurons.

**Figure 7:**
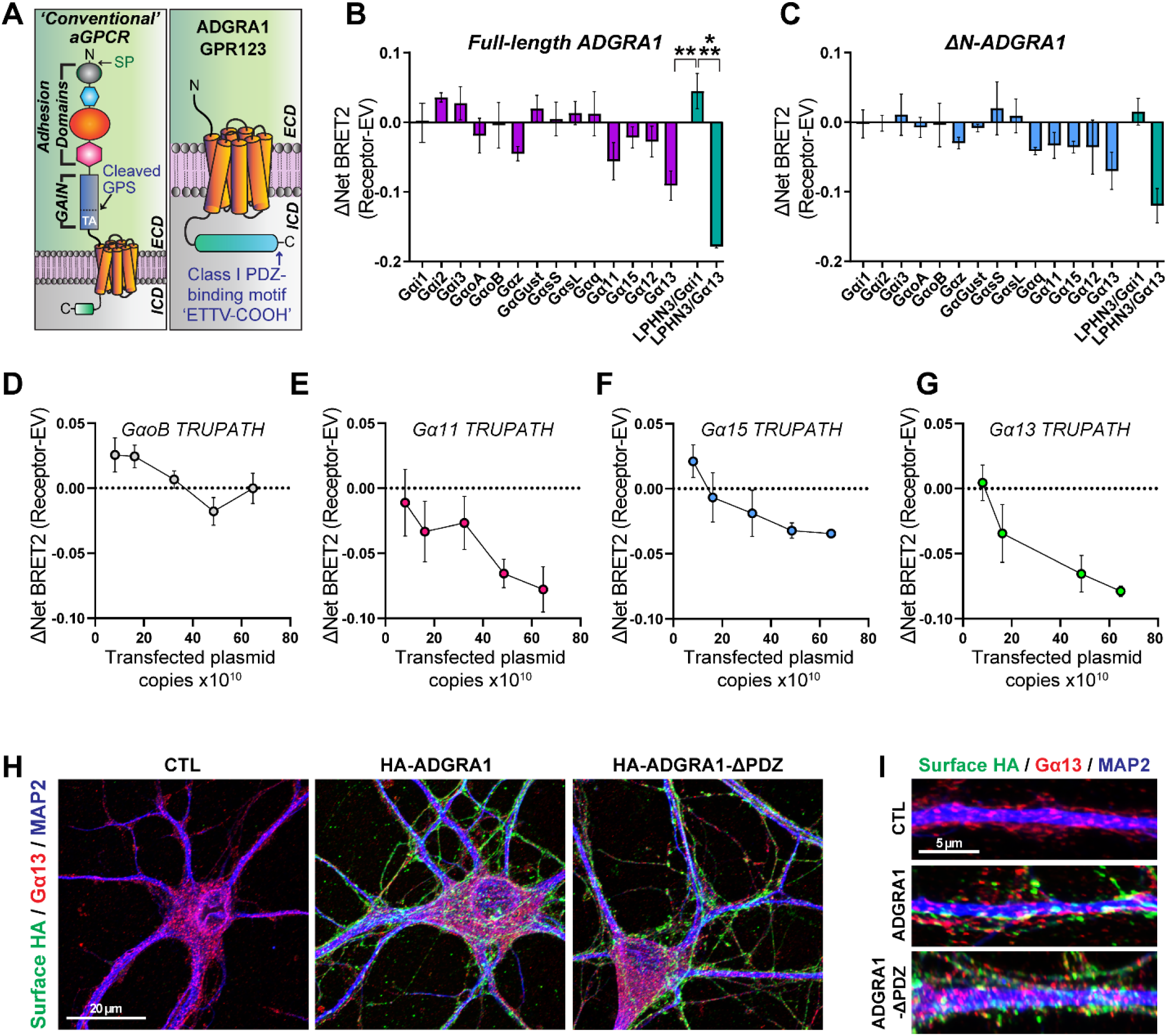
ADGRA1 engages several G proteins including Gα13. **A,** diagram of ADGRA1 protein domains compared to standard aGPCRs. **B,** TRUPATH BRET2 analysis of full-length ADGRA1. A LPHN3 construct with either the Gαi1 or Gα13 TRUPATH sensors were used as negative or positive controls, respectively. **C,** similar to **B**, except using an ADGRA1 mutant lacking the short N-terminal extracellular sequence prior to the 7-TM GPCR. **D-G,** ADGRA1 plasmid dose-response curves using indicated TRUPATH BRET2 biosensors. **D,** ADGRA1 plasmid dose-response experiments for GαoB. **E,** similar to **D**, except for Gα11. **F,** similar to **D**, except for Gα15. **G,** similar to **D**, except for Gα13. **H,** representative immunocytochemistry for surface HA-ADGRA1 and intracellular Gα13 and MAP2 in indicated conditions. **I,** representative MAP2-labeled dendrite co-labeled for surface HA and intracellular Gα13. Numerical data are means ± SEM. See Figure S12 for additional characterization of ADGRA1 constructs and quantification of ADGRA1/Gα13 co-localization. Statistical significance was determined via one-way ANOVA with *post hoc* Tukey test (**, p<0.01; ***, p<0.001).

## Discussion

The cell surface receptors and signaling pathways that establish inhibitory circuitry remain poorly understood. Our studies identify ADGRA1 as a synaptic receptor highly enriched in hippocampal interneurons. ADGRA1 functions as a critical receptor within interneurons and controls inhibitory synaptic strength in the DG circuit. Furthermore, ADGRA1 activates several G proteins including Gα13. Recent studies found that the Gα13 signaling pathway is involved in establishing hippocampal PV inhibitory circuitry^29^. We postulate that ADGRA1 is localized at synapses in PV and SST interneurons, where it controls a Gα13-dependent pathway that establishes inhibitory circuits in a cell type-specific manner. Our results suggest that ADGRA1 controls the function of PV synapses rather than their overall density in the DG circuit. Thus, ADGRA1 likely functions in mature hippocampal inhibitory circuits rather than during development.

The endogenous ligands for ADGRA1 are unknown. Contrary to all other aGPCRs, ADGRA1 lacks a GAIN domain and therefore cannot utilize the primary modes of aGPCR activation mechanisms previously studied involving either the TA or conformational coupling of the NTF and CTF^30^. We found that the short extracellular region of ADGRA1 is not involved in basal G protein activation. ADGRA1 may be activated by small molecule ligands similar to other GPCRs. For example, recent studies identified that the aGPCR GPR97 is activated by glucocorticoids^31^. ADGRA1 may use similar steroid ligands independent from the GAIN and TA. Alternatively, ADGRA1 may form heterodimers with other GPCRs which are necessary for signaling function. Yet another potential mechanism involves ADGRA1 acting as a constitutively active receptor which is regulated instead through spatial restriction of its localization. Future studies identifying how ADGRA1 is activated will advance our understanding of both inhibitory circuit assembly and aGPCR activation mechanisms.

While ADGRA1 lacks an extensive extracellular region, it contains a large intracellular C-terminal tail with a PDZ-binding motif. This elaborate tail region is likely involved in ADGRA1 synaptic function. Other aGPCRs, including Latrophilins (Lphns), exhibit extensive C-terminal tails essential for their functions. For example, the intracellular tail of Lphn serves as a scaffold for several protein-protein interactions, including postsynaptic excitatory SHANK proteins^32,33^. Lphn3 mutants lacking the C-terminal tail are incapable of rescuing Lphn3-dependent synaptic deficits^9^. The large intracellular tail of ADGRA1 likely also functions as a scaffold for several protein-protein interactions, and may be important for ADGRA1 subcellular localization. We found that deleting the C-terminal PDZ-binding motif had no effect on ADGRA1 synaptic localization. Thus, other sequence features in the large C-terminal tail are likely responsible for the subcellular targeting of ADGRA1. Interestingly, the C-terminal tail of closely related ADGRA3 has been shown to interact with DLG1/SAP97, a scaffold in the MAGUK family that contains three PDZ domains^34^. Thus, the ADGRA intracellular tail may function as a scaffold for multiple protein-protein interactions that cluster at the synapse.

We were unable to definitively determine the subcellular localization of endogenous ADGRA1 due to lack of a reliable antibody. Future studies will be required to determine its endogenous localization and protein-protein interaction partners. Moreover, while we observed localization of ADGRA1 at synapses, consistent with previous studies^21^, we were unable to definitively measure pre- or postsynaptic localization due to the resolution limits of confocal microscopy. Super-resolution methods will be required to determine the precise sub-synaptic position of ADGRA1. While we determined that ADGRA1 functions in both PV and SST interneurons, the molecular mechanisms of these functions remain unclear. Moreover, our mRNA expression analysis shows that Adgra1 is expressed in various other brain regions, including cortical layer 6 and the thalamus. ADGRA1 may engage distinct signal transduction pathways within these different cell types, or may interact with different sets of protein-protein interactions within these cells to direct the specificity of PV vs. SST synaptic function.

Collectively, these studies identify an orphan receptor selective to PV and SST hippocampal interneurons which is essential for their synaptic function. Further knowledge of the cell surface receptors and downstream signaling pathways responsible for synapse specific circuit establishment will increase our understanding of how remarkably diverse synaptic connections process information and generate behaviors.

## Methods

### RESOURCES TABLE

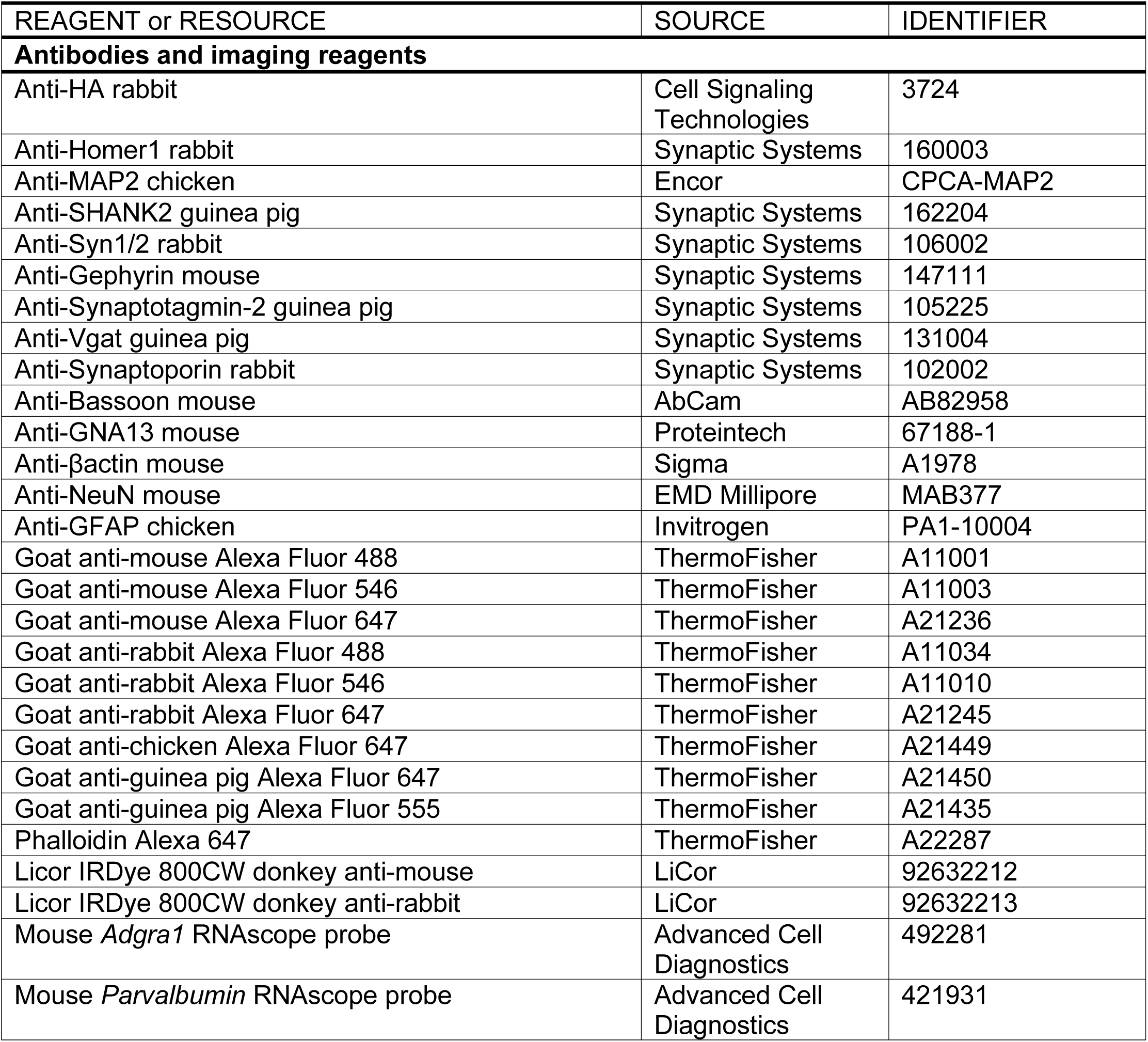
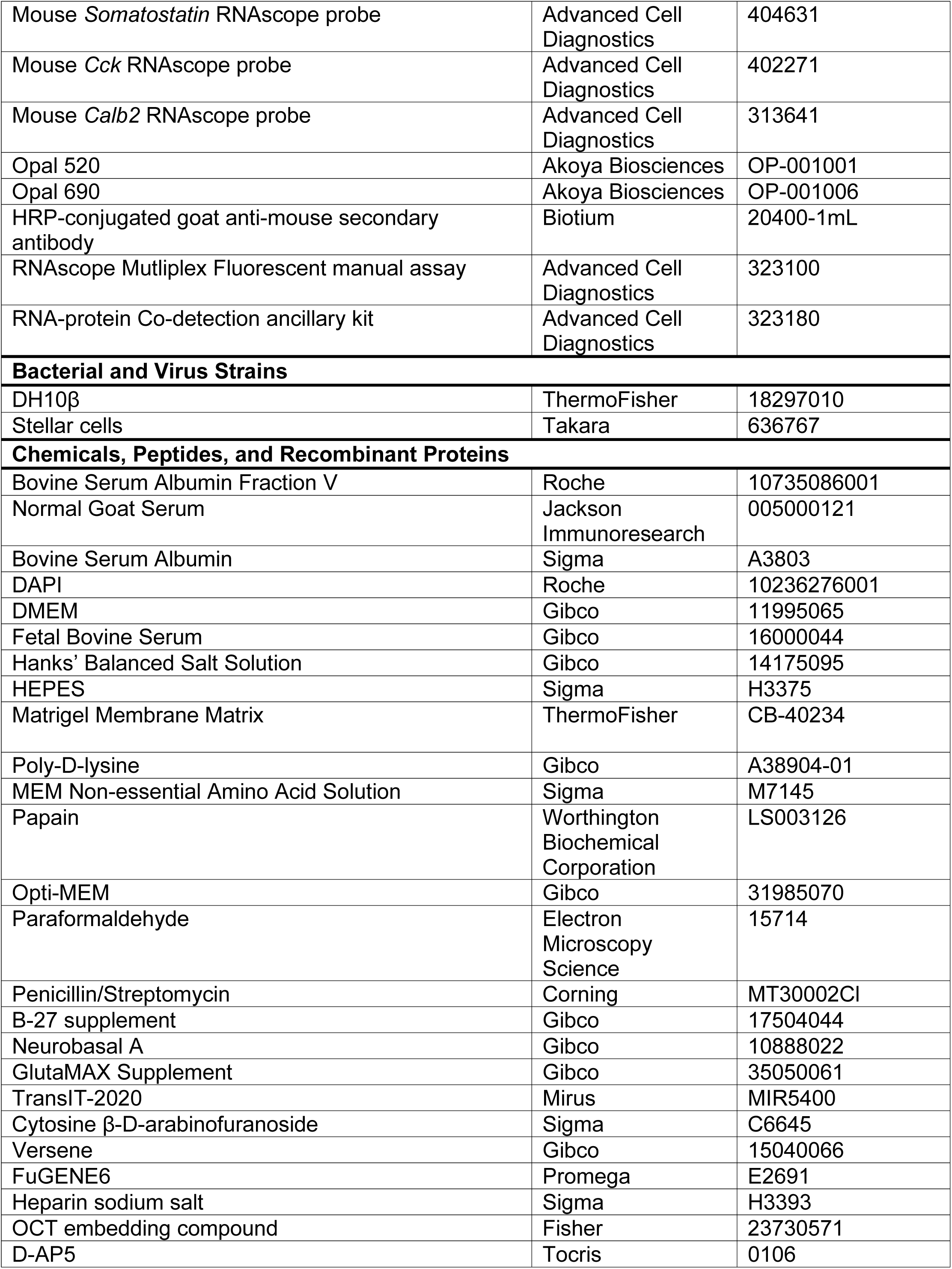
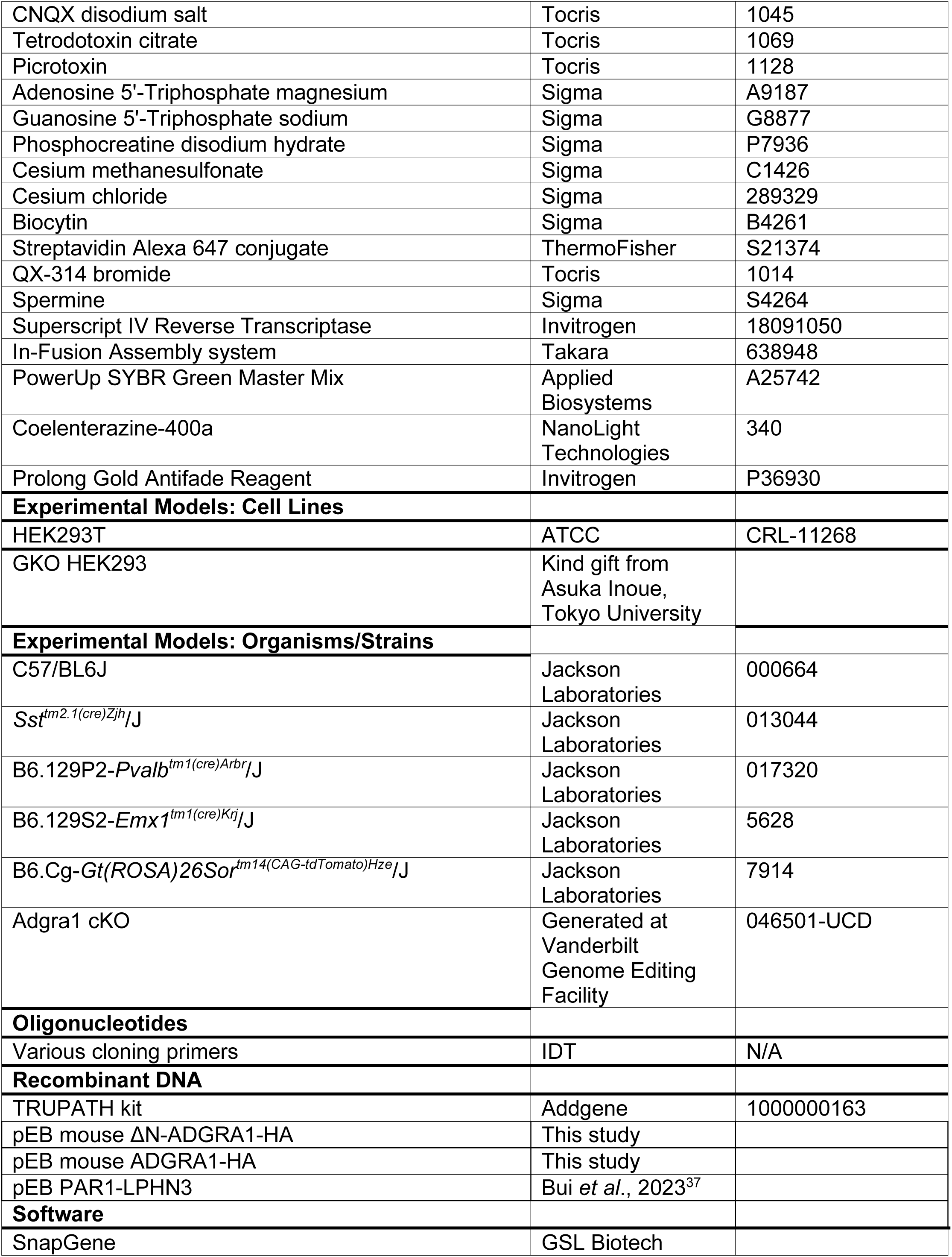
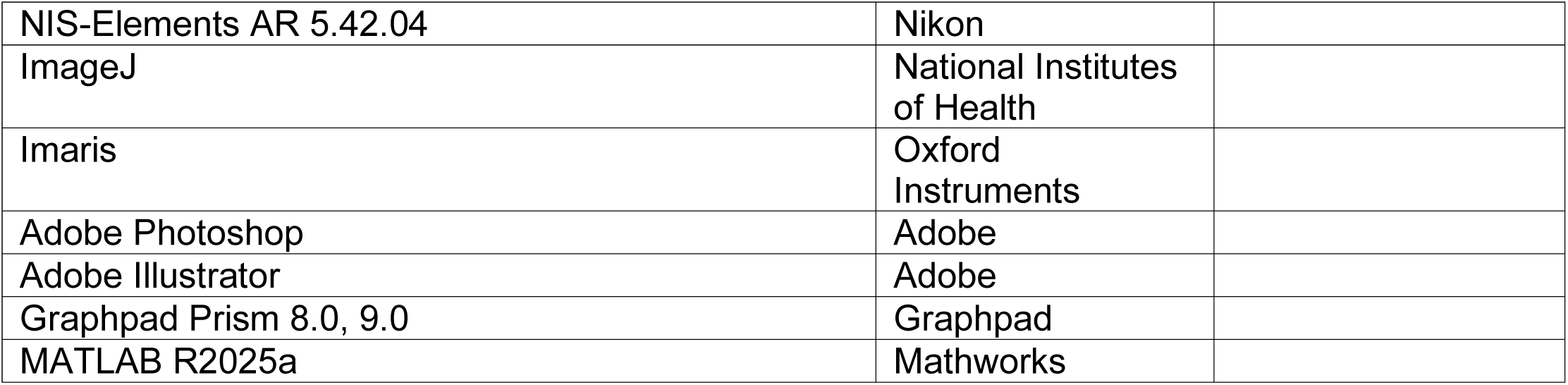

### RESOURCE AVAILABILITY

#### Lead contact

Further information and requests for resources and reagents should be directed to and will be fulfilled by the lead contact, Richard C. Sando.

#### Materials availability

All materials generated in this study will be openly shared upon request.

#### Data and code availability

No original code has been generated in this study.

### EXPERIMENTAL MODELS AND SUBJECT DETAILS

#### Mice

Mice were weaned at 18-21 days of age and housed in groups of 2 to 5 on a 12 h light/dark cycle with food and water *ad libitum*. Vanderbilt Animal Housing Facility: All procedures conformed to National Institutes of Health Guidelines for the Care and Use of Laboratory Mice and were approved by the Vanderbilt University Administrative Panel on Laboratory Animal Care. Primary hippocampal cultures were generated from P0 pups from Adgra1 cKO and C57BL/6J mice (Jax, Cat#000664). For generation of Adgra1 cKO mice, sperm was obtained from KOMP MMRRC (Stock number 046501-UCD), and the line was generated at the Vanderbilt Genome Editing Facility. C57BL/6J female mice (Jax, Cat#664) were superovulated for *in vitro* fertilization using the CARD method (Takeo *et al.*, 2019, Methods Mol Biol^38^). There was a 52% fertilization rate, and 100 embryos were transferred generating 25 pups. The following CRE driver lines and reporter lines were used: EMX-Cre, B6.129S2-*Emx1^tm1(cre)Krj^*/J (Jax, Cat#5628); PV-Cre, B6.129P2-*Pvalb^tm1(cre)Arbr^*/J (Jax #017320); SST-Cre, *Sst^tm2.1(cre)Zjh^*/J (Jax, Cat#013044); Ai14, B6.Cg-*Gt(ROSA)26Sor^tm14(CAG-tdTomato)Hze^*/J (Jax #007914).

#### Cell Lines

GKO HEK293 cells used for all TRUPATH BRET2 assays were originally a kind gift from Asuka Inoue (Tokyo University, Japan), and were provided to our studies as a kind gift from Drs. Vsevolod Gurevich and Chen Zheng (Vanderbilt University). GKO HEK293 cells were maintained in DMEM (Gibco, Cat#11995065) containing 10% FBS (Gibco, Cat#16000044), 1X Penicillin-Streptomycin (Corning, Cat# MT30002Cl), and 1x MEM Non-essential Amino Acid (NeA) Solution (Sigma, Cat# M7145) at 37°C and 5% CO_2_. HEK293T cells (ATCC # CRL-11268 were maintained in DMEM (Gibco, Cat#11995065) containing 10% FBS (Gibco, Cat#16000044), 1X Penicillin-Streptomycin (Corning, Cat#MT30002Cl) at 37°C and 5% CO_2_. Cell lines were maintained for a maximum of 25 passages.

#### Primary hippocampal cultures

For immunocytochemistry, cover glasses (#0, 12 mm, Carolina Biological Supply Company, Cat#633009) were placed into 24-well plates (Genesee, Cat#25-107MP) and coated for 2 hrs with 100 µL of 50 µg/mL poly-D-lysine (Gibco, Cat#A38904-01) in a 37°C tissue culture incubator. Excess poly-D-lysine was removed, coverslips were washed 3x with sterile dH_2_O and then dried for 30 mins. For primary cells, bilateral hippocampi were dissected from P0 mouse pups and collected tissue was dissociated by papain (Worthington Biochemical Corporation, Cat#LS003126) digestion for 20 min at 37°C. After the digestion, the excess papain solution was removed and tissue was washed twice with plating media containing: 5% fetal bovine serum (Life Technologies, Cat#16000044), B27 (Gibco, Cat#17504044), 1:50), 0.4% glucose, and 2 mM glutamine in 1x MEM. Immediately after the second wash, tissue was triturated using a 1000 µL pipette. Next, cells were filtered through a 70 µm cell strainer (Corning, Cat#431751), and plated at a density of 40,000 cells per dish/well in 1 mL plating media. Culture media was exchanged 24 hrs later (at DIV1) to growth medium, which contained 5% fetal bovine serum, B27, 2 mM glutamine in Neurobasal A (Gibco, Cat#10888022). Cytosine β-D-arabinofuranoside (Sigma, Cat#C6645) was added to a final concentration of 4 µM on DIV3 along with a 50% growth media exchange. Primary hippocampal cultures were infected with respective lentiviral conditions at DIV1, and staining experiments were conducted between DIV10-14.

#### Plasmids

Mouse ADGRA1 overexpression cDNAs used in HEK293T cell experiments were encoded in the pEB Multi-Neo vector (Wako Chemicals, Japan), and empty vector controls were empty pEB Multi-Neo. Mouse ADGRA1 corresponded to Uniprot #Q8C4G9 and contained a N-terminal HA tag for expression studies. ΔN-ADGRA1 contained the first 22 residues of ADGRA1 (MTQWDLKTVLSLPQYPGEFLHP) replaced with an 8x glycine flexible sequence. Lentiviral NLS-GFP-Cre or NLS-GFP-ΔCre were encoded in a 3^rd^ generation lentiviral backbone and driven by the EF1α promoter. All molecular cloning was conducted with the In-Fusion Assembly system (Takara, Cat#638948).

#### Antibodies

The following antibodies and reagents were used at the indicated concentrations (IHC-immunohistochemistry, ICC-immunocytochemistry, IB-immunoblot): anti-HA rabbit (Cell Signaling Technologies, Cat# 3724, 1:1,000, ICC, 1:2,000 IB), anti-Homer1 rabbit (Synaptic Systems, Cat#160003, 1:500 IHC), anti-Bassoon mouse (AbCam, Cat#ab82958, 1:500 IHC,1:1,000 ICC), anti-SHANK2 guinea pig (Synaptic Systems, Cat#162204, 1:1,000 ICC, 1:500 IHC), anti-Syn1/2 rabbit (Synaptic Systems, Cat#106002, 1:500 IHC), anti-VGAT guinea pig (Synaptic Systems, Cat#131004, 1:500 IHC), anti-Synaptoporin rabbit (Synaptic Systems, Cat#102002, 1:500 IHC), anti-Gephyrin mouse (1:2,000 ICC, Synaptic Systems, Cat#147111) anti-Parvalbumin rabbit (SWANT, Cat#PV27a, 1:2,000 IHC), anti-Synaptotagmin-2 (Synaptic Systems, Cat#105225,1:500 IHC), Anti-βactin mouse (Sigma, Cat#A1978, 1:10,000 IB) and corresponding fluorescently-conjugated goat secondary antibodies from Life Technologies (1:1,000).

#### TRUPATH BRET2

HEK G protein K.O cells were plated into 12-well plates at a density of 3-4 x 10^5^ cells in 1 mL per well. HEK G K.O. media contained 1x DMEM (Gibco, Cat# 11995065) plus 10% FBS (Gibco, Cat#16000044), 1X Penicillin-Streptomycin (Corning, Cat#MT30002Cl) with 1x MEM Non-essential Amino Acid (NeA) Solution (Sigma, Cat# M7145). Cells were co-transfected 16-24 hrs after plating with receptor-of-interest and TRUPATH plasmids at 1:1:1:1 DNA ratio (receptor:Gα-RLuc8:Gβ:Gγ-GFP2) via TransIT-2020 (Mirus, Cat# MIR5400). Each condition required 97 µL of room-temperature 1x Opti-MEM (Gibco, Cat# 31985070), 1 µL each DNA plasmid at 1 µg/µL concentration), and 3 µL of room-temperature and gently vortexed TransIT-2020 reagent. The TransIT-2020:DNA complexes mixtures were gently mixed via pipetting 10 times and incubated at room temperature for 20 mins before adding dropwise in the well. The plate was rocked gently side to side and incubated at 37°C for 24 hours before harvesting. In each well, media was aspirated, and cells were washed with 1 mL warm PBS. Cells were detached with 300 µL warm Versene (Gibco, Cat# 15040066) and incubated at 37°C for 5 mins then resuspended via gentle pipetting 10 times. Cells were plated in complete DMEM containing 1x NeA at 200 µL with a density of 30,000–50,000 cells per well in Matrigel-coated 96-well assay plate. Each experimental condition was plated into three separate wells within the 96-well assay plate. BRET2 assays were performed 48 hrs after transfection. In each well, media was aspirated, and cells were incubated in 80 µL of 1x Hanks’ balanced Salt Solution (Gibco, Cat# 14175095) with 20 mM HEPES (Sigma, Cat# H3375, pH 7.4) and 10 µL 100 µM Coelenterazine-400a (NanoLight Technologies, Cat# 340) diluted in PBS. After 10 mins of incubation, BRET2 intensities were measured using a BERTHOLD TriStar^2^ LB 942 Multimode Reader with Deep Blue C filter (410nm) and GFP2 filter (515 nm). The BRET2 ratio was obtained by calculating the ratio of GFP2 signal to Deep Blue C signal per well. The BRET2 ratio of the three wells per condition were then averaged. Net BRET2 was subsequently calculated by subtracting the BRET2 ratio of cells expressing donor only (Gα-RLuc8) from the BRET2 ratio of each respective experimental condition. Net BRET2 differences were then compared as described in the Figures, by subtracting Net BRET2 ratios of conditions overexpressing indicated receptors compared to empty vector (EV). For plasmid copy-dependent BRET2 experiments, conditions were transfected in a 12-well plate format with the same total amount of plasmid DNA and varying copies of experimental plasmid, adjusted to the same total amount with empty vector (pEB-multi).

#### Ribotag immunoprecipitations

Hippocampal tissue was dissected as quickly as possible on a foil wrapped metal block on ice, gently transferred to a microcentrifuge tube and flash frozen in liquid nitrogen. Tissue was then stored at -80°C until use in the RiboTag procedure. The benchtop surface and all supplies were sprayed with RNAse zap (Invitrogen, Cat#AM9780). The Homogenization Buffer (HB), Supplemented Homogenization Buffer (HB-S), and Lysis Buffer (LB) solutions were prepared as follows. HB: 1% IGEpal (Spectrum Chemical MFG Corp, Cat#I1112), 100 mM KCl (Sigma, Cat#P5405-500g), 50 mM Tris, pH 7.4 (Sigma, Cat#T6066-500G), and 12 mM MgCl_2_ (Sigma-Aldrich, Cat#63069-100mL) to final volume with nuclease-free dH_2_O. HB-S (for a 10 mL solution): 9.424 mL previously prepared HB, 0.2 mL of Cycloheximide (5 mg/mL, Sigma, Cat#C7698-1G), 0.1 mL Mammalian Protease Inhibitor Cocktail (Sigma-Aldrich, Cat#P8340-1mL), 0.2 mL Heparin (50 mg/mL, sodium salt from porcine intestine, (Sigma, Cat#H3393-100KU), 0.01 mL of RNAsin (Promega, Cat#N251A), 0.01 mL of 1M DTT (Sigma, Cat#D0632-10G). LB: 1.2 mL of RLT Plus Buffer (Qiagen, Cat#1030963), 0.012 mL of β-Mercaptoethanol (Sigma, Cat#M3148-100mL). Once solutions were prepared, tissue was weighed and homogenized in 10% weight/volume HB-S in a dounce homogenizer (Kimble, Cat#885301-0007). The homogenization was done by homogenizing each sample 20 times with the pestle, taking care that as few bubbles as possible were formed. Once homogenized, the homogenate was transferred to fresh microcentrifuge tubes and spun in a cooled centrifuge for 10 mins at 4°C and 10,000 x g. The supernatant was collected, and 50 µL of supernatant was transferred into a new tube and combined with 350 µL of the previously mentioned LB to generate the Input sample, which was flash frozen in liquid nitrogen. During the centrifugation Pierce Anti-HA magnetic beads (Thermo Scientific, Cat#88836) were prepared as follows. 30 µL of beads per sample were placed into a microcentrifuge tube in a magnetic rack and the storage buffer was removed. Beads were subsequently washed in 1000 µL of HB-S on a gentle rotator for 10 mins at 4°C. Beads were collected, the wash was removed and 30 µL of HB-S per sample were added to the beads. Once the beads were resuspended, the remainder of the tissue homogenate supernatant was added and incubated in dark on a gentle rotator at 4°C overnight. After the overnight incubation, beads were then washed 3 times for 10 mins in High Salt Buffer (HSB) (for a 10 mL solution): 5.18 mL Nuclease Free Water, 3.0 mL of 1M KCl, 1.0 mL 10% IGEpal, 0.5 mL of 1.0 M Tris, pH 7.4, and 0.12 mL of 1M MgCl_2_, 0.2 mL of Cycloheximide (5 mg/mL), 0.01 mL of 1M DTT. The supernatant was discarded and immediately after the final wash, 350 µL of LB was added to the beads and the samples were resuspended on a vortex mixer on speed 4 for 30 seconds. Then the vortexed samples were placed back on the magnetic rack and the supernatant was drawn off and placed in a fresh Eppendorf. Then the input samples frozen from the previous day were vortexed on speed 4 for 30 seconds. RNA was subsequently purified from Input and IP samples using the RNeasy Micro Kit (Qiagen, #74134), eluting in 20 µL of RNase-free water. The purified eluted RNA was converted to cDNA using SuperScript IV (Invitrogen, Cat#18090050) with random hexamers.

#### RT-qPCR

RNA was extracted from primary hippocampal neurons between DIV10-12. Each well was briefly washed with 1x PBS, followed by the addition of 250 µL Trizol. The plate was rocked for 5 mins at high setting at RT, and all samples were pooled into a single Eppendorf tube, bringing the final volume to 1 mL with additional Trizol. The mixture was incubated at room temperature for 5 mins, then 200 µL chloroform was added and mixed gently by inverting the tube 20 times. After 3-min incubation at RT, samples were centrifuged at 12,000 x g for 15 mins at 4°C. The upper aqueous phase (∼500 µL) was carefully collected without disturbing the interphase and transferred to a new tube. An equal volume (500 µL) of 2-propanol was added and mixed by inversion 20 times, followed by a 10-min incubation at RT. The RNA was precipitated by centrifugation at 12,000 x g for 10 mins at 4°C. The supernatant was discarded, and the pellet was washed two times with 500 µL of 70% ethanol by spinning at 7,000 x g for 5 mins at 4°C. After removing all remaining supernatant, the pellet was immediately resuspended in 100 µL of RNase-free dH₂O pre-warmed to 55°C, then incubated at 55°C for 10 minutes. During this incubation, Buffer RLT with β-Mercaptoethanol (10 µL per 1 mL RLT) was made. Finally, the tube was gently tapped to fully dissolve the RNA pellet, and 350 µL of the RLT/β-Mercaptoethanol solution was added to the sample. Total RNA was isolated using the RNeasy Micro Kit (Qiagen, Cat#74134), and then used for cDNA synthesis with Super Script IV (Invitrogen, Cat#18090050) with random hexamers. qPCR with PowerUp SYBR Green Master Mix (Applied Biosystems, Cat#A25742) was used for quantification of relative gene expression and subsequently normalized to the expression of GAPDH. qPCR primer pairs (PrimeTime, IDT) were used mentioned in Table 1.

**Table 1.**
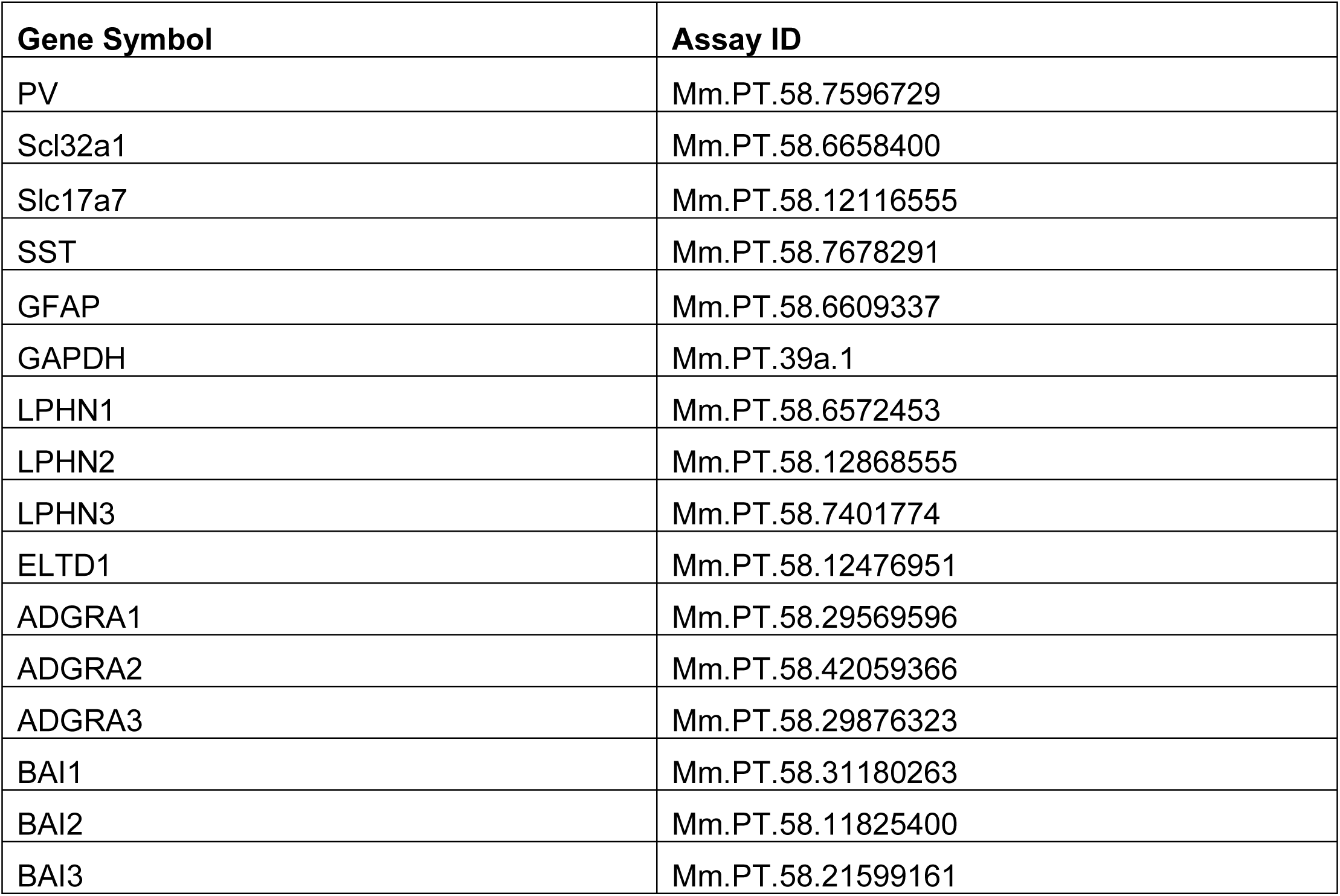
RT-qPCR primers.

#### Surface labeling and immunocytochemistry

Cells were washed briefly once with PBS, fixed with 4% PFA (Electron Microscopy Sciences, Cat#15714)/4% sucrose/PBS for 20 min at 4°C, and washed 3 x 5 min in PBS. After the final wash, blocking buffer containing 4% BSA (Sigma, Cat#10735086001)/3% normal goat serum (Jackson Immunoresearch, Cat#005000121)/PBS was applied for 30 mins. For surface immunolabeling of HA-ADGRA1, samples were subsequently incubated overnight at 4°C with anti-HA rabbit (Cell Signaling Technologies, Cat#3724) primary antibody solution diluted in blocking buffer. The next day, samples were washed 3 times with PBS, followed by permeabilization with 0.2% Triton X-100/PBS for 5 min at room temperature. After permeabilization, samples were transferred into blocking buffer and incubated at RT for 30 min. For total labeling, following primary antibodies were used: anti-MAP2 chicken (1:5,000; EnCor, Cat#CPCA-MAP2), anti-SHANK2 guinea pig (1:1,000; Synaptic Systems, Cat#162204), anti-Bassoon mouse (1:1000; AbCam, Cat#AB82958), anti-GNA13 mouse (1:1000; Proteintech, Cat#67188-1), and anti-Gephyrin mouse (1:2,000; Synaptic Systems, Cat#147111). Samples were incubated with primary antibodies for 2 hr at RT. Samples were then washed 5 x 5 min in PBS, incubated with fluorescently conjugated secondary antibodies diluted in blocking buffer for 1 hr at room temperature. Corresponding secondary antibodies were goat anti-chicken 647 (1:1,000; ThermoFisher, Cat#A21449), goat anti-rabbit 488 (1:1,000 ThermoFisher, Cat# A11034), goat anti-guinea pig 555 (1:1,000; ThermoFisher, Cat#A21435), and goat anti-mouse 546 (1:1,000; ThermoFisher, Cat#A11003). Samples were labeled with DAPI (Sigma, Cat#10236276001) diluted into PBS for 5 min, then washed 4 x 5 mins with PBS. Samples were mounted on UltraClear microscope slides (Denville Scientific, Cat# M1021) using 10 µL ProLong Gold antifade reagent (Invitrogen, Cat#P36930) per coverslip.

#### Immunoblotting

Primary hippocampal cultures were briefly washed 1X with PBS, and samples were collected in sample buffer containing 312.5 mM Tris-HCl pH 6.8, 10% SDS, 50% glycerol, 12.5% 2-mercaptoethanol, bromophenol blue, and protease inhibitors (Roche, Cat# 11873580001) and run on 12% SDS-PAGE gels (BioRad mini-protean TGX, Cat#4561024) at 30 mA/gel constant current. The HiMark Prestained Protein Standard (Invitrogen, Cat# LC5699) was used as a protein molecular weight ladder. Protein was transferred onto nitrocellulose transfer membrane in transfer buffer (25.1 mM Tris, 192 mM glycine, 20% methanol) at 200 mA constant current for 2 hrs at 4°C. Membranes were blocked in 4% bovine serum albumin (BSA, Sigma, Cat#10735086001)/TBST (20 mM Tris-HCl pH 7.4, 150 mM NaCl, 0.05% Tween-20) for 1 hr at room temperature, incubated in primary antibodies diluted into 4% BSA/TBST overnight at 4°C (anti-HA rabbit 1:2,000; anti-βactin mouse 1:10,000), washed 3 x 5 mins in TBST, incubated in corresponding secondary antibodies (Licor IRDye 800CW donkey anti-mouse, Cat#92632212; anti-rabbit #92632213) diluted 1:10,000 into TBST, washed 5 x 5 mins in TBST, and imaged on a Licor Odyssey system. Immunoblot intensities were measured using LiCor Image Studio.

#### Immunohistochemistry

Mice were briefly anesthetized with isoflurane and transcardially perfused with 10 mL room temperature heparinized (10 U/mL, Sigma, Cat#H3393) PBS, followed by 25 mL room temperature 4% PFA/PBS. The brains were post-fixed 2 hrs at 4°C, washed with PBS, cryoprotected in gradients of 10% sucrose/PBS, 20% sucrose/PBS, and 30% sucrose/PBS, rapidly embedded in OCT (Fisher, Cat#23730571), and sliced on a cryostat at 25 µm. The 25 µm thick, free-floating sections were washed 1 x 5 min with PBS/0.1% Triton X-100, blocked for 1 hr at room temperature in blocking buffer containing 4% BSA (Sigma Cat#10735086001)/3% normal goat serum (Jackson Immunoresearch Cat#005000121)/0.1% Triton-X100/PBS and incubated overnight at 4°C with primary antibodies diluted in blocking solution. The following primary antibodies were used: anti-vGAT guinea pig (1:1,000; Synaptic Systems Cat#131004), anti-Bassoon mouse (1:500; AbCam, Cat#AB82958), anti-Synaptoporin rabbit (1:500; Synaptic Systems, Cat#102002), anti Syn1/2 rabbit (1:500; Synaptic Systems Cat#106002), Anti-Synaptotagmin-2 guinea pig (1:500; Synaptic Systems, Cat#105225), anti-Homer1 rabbit (1:500; Synaptic Systems, Cat#160003), anti-Parvalbumin rabbit (1:2,000; Swant, Cat#PV27a), and anti-SHANK2 guinea pig (1:500; Synaptic Systems, Cat#162204). Primary antibody incubation was followed by 3 x 5 min washes in PBS/0.1% Triton X-100 and 2 hr room temperature incubation with corresponding fluorescently labeled secondary antibodies in blocking buffer. Samples were labeled with DAPI (Sigma, Cat#10236276001) diluted into PBS/0.1% Triton X-100 for 15 mins, washed 5 x 5 mins with PBS/0.1% Triton X-100, and mounted on glass slides coated in 0.1% Triton-X100/PBS, dried briefly, and covered with ProLong Gold antifade reagent (Invitrogen, Cat#P36930) and cover glass (Corning, Cat#2980-246).

#### Lentivirus production for culture experiments

Lentiviruses were packaged in HEK293T cells from ATCC (CRL-11268). For lentiviral production, co-transfection of the expression shuttle vector and the three helper plasmids (pRSV-REV, pMDLg/pRRE and vesicular stomatitis virus G protein (VSVG)) was done with FuGENE6 (Promega E2691) using 2.5 µg of each plasmid per 9.6 cm^2^. Lentiviral-containing medium was collected 48 hrs after transfection, briefly spun down 5,000 xg for 5 mins for removal of cellular debris and then stored at 4°C. The LV genomic titer was estimated using PowerUp™ SYBR™ Green Master Mix for qPCR (Applied Biosystems, A25742) with the following primers: F - ccactgctgtgccttggaatgc, and R - aatttctctgtcccactccatccag. Shuttle plasmids at 10x serial dilutions (1×10^5^ - 1×10^9^ copies/mL) were used for generating a standard curve. After quantification, the LVs were directly applied to primary neuron culture medium.

#### RNA in situ hybridizations

Wild-type C57BL/6J (Jackson, Cat#000664) mice were taken from their home cages at postnatal day (P) 5, P10, P21, and P30, and whole brain tissue was collected in the following manner. Brains were rapidly dissected following brief anesthesia with either ice (P5) or isoflurane (P10, P21, P30) and placed in a rectangular cryomold (Epredia Peel-a-way, Cat#18-30) which was flash-frozen in liquid N_2_ for 15 seconds to allow for indirect exposure of the tissue with liquid N_2_. The brain was subsequently embedded in O.C.T. Compound (Fisher Healthcare, Cat#4585) within a second cryomold using a bath of 2-methylbutane (Sigma-Aldrich, Cat#M32631-46) chilled with dry ice. Once frozen, the blocks were stored at -80°C until cryosectioning. The frozen blocks were removed from the -80°C freezer and allowed to equilibrate in the cryostat (Leica CM 1950, Cat#047742456) at -20°C for 1 hr. The blade for slicing (Sakura, Cat#4689), forceps, razor blade for block trimming, and paintbrushes for manipulating sections, and the anti-roll plate (Leica, Cat#14047742497) were also all placed in the cryostat and allowed to equilibrate. Tissue was sectioned at 15 µm and mounted directly onto room temperature Diamond White Glass microscope slides (Globe Scientific Inc., white frosted 25×75×1mm, charged +/+, Cat#1358W). Once mounted, the slides were kept in the cryostat until all sectioning was complete. Sections were subsequently dried at -20°C for 1 hr, then stored at -80°C.

Tissue Pre-Treatment: The RNAscope Multiplex Fluorescent manual assay (Advanced Cell Diagnostics, Cat#323100) was carried out using the Fresh Frozen sample preparation according to the manufacturer’s protocol as described below. The RNAscope Hydrogen Peroxide (Advanced Cell Diagnostics, Cat#322335) and RNAscope Protease IV (Advanced Cell Diagnostics, Cat#322336) reagents were set out on the benchtop to equilibrate to room temperature. Sections were removed from the -80°C freezer and placed immediately into ice cold 4% PFA (Electron Microscopy Sciences, Cat#15714)/PBS (MP Biomedicals, Cat#2810306) within a glass slide holder (Epredia RA Lamb Glass Coplin Jar, Fisher, Cat#E94). Slides were incubated at 4°C for 15 mins to fix the tissue and subsequently washed twice with 1x PBS. Slides were subsequently dehydrated in the following ethanol (Decon Laboratories, Inc., 200 proof, Cat#2705HC) gradient: 50% EtOH/ddH_2_O for 5 minutes, 70% EtOH/ddH_2_O for 5 mins, followed by two treatments with 100% EtOH (50 mL of each treatment). After the final 100% step, the slides were placed section side up on a paper towel and allowed to dry for 5 mins. Then a hydrophobic pen (IHC World, super pap pen, Cat#SPR0905) was used to draw a barrier around each section, which dried at room temperature for 5 mins. While the barriers were drying, the HybEZ™ Humidity Control Tray with lid (Advanced Cell Diagnostics, Cat#310012) was prepared. A sheet of HybEZ™ Humidifying Paper (Advanced Cell Diagnostics, Cat#310025) was placed on the bottom of the tray and sprayed with ddH_2_O until damp. The EZ-Batch™ Slide Holder (Advanced Cell Diagnostics, Cat#310017) was placed inside the humidity control tray and the slides were placed in the holder. Three drops of hydrogen peroxide were added to each section. The cover was placed over the humidity control tray and the slides were left to incubate for 10 mins at room temperature. Once the incubation was complete, the slides were washed twice with ddH_2_O, removing excess liquid after each wash with a vacuum aspirator. Slides were reinserted into the slide holder and 4 drops of Protease IV were added to each section, followed by incubation at room temperature for 30 mins. While the slides were incubating the RNAscope Probes were prepared in the following manner. Probes (Adgra1-C3 #492281, PV-C2 #421931, SST-C2 #404631, CCK-C2 #402271, Calb2-C2 #313641) were placed in a heat block at 40°C for 10 mins, and subsequently removed from the heat and incubated at room temperature for 10 mins. The probes were combined to form a probe mix consisting of 100 µl of probe diluent per section, and 2 µL each of the C2 and C3 probes per section, respectively. The HybEZ™ II Oven (Advanced Cell Diagnostics, Cat#321720) was then pre-warmed to 40°C. Once the Protease IV incubation was complete, the slides were washed twice with 1x PBS, placed back in the slide holder and humidity control tray and 100 µL of probe mix were added to each section. The tray was placed in the HybEZ™ Oven for 2 hrs at 40°C to hybridize the probes. While the incubation was occurring, the 1x RNAscope wash buffer solution (Advanced Cell Diagnostics, #310091) and 5x Saline Sodium Citrate (SSC) buffer were prepared. The 20x SSC stock contained 175.3 g of NaCl (Fisher Chemical, certified ACS, crystalline, #S271-1) and 88.2 g of sodium citrate (Fisher Chemical, dihydrate, granular, Cat#S279-500) in ddH_2_O, pH 7.0. Following the 2-hr period, slides were washed twice with 1x RNAscope wash buffer and placed in 5x SSC buffer overnight at 4°C.

RNAscope Multiplex Fluorescent Assay: The following reagents were equilibrated at room temperature for 1 hr: RNAscope Multiplex FL v2 AMP1 (Advanced Cell Diagnostics, 323101), RNAscope Multiplex FL v2 AMP2 (Advanced Cell Diagnostics, 323102), RNAscope Multiplex FL v2 AMP3 (Advanced Cell Diagnostics, 323103), RNAscope Multiplex FL v2 HRP C1 (Advanced Cell Diagnostics, 323104), RNAscope Multiplex FL v2 HRP C2 (Advanced Cell Diagnostics, 323105), RNAscope Multiplex FL v2 HRP C3 (Advanced Cell Diagnostics, 323106) and RNAscope Multiplex FL v2 HRP Blocker (Advanced Cell Diagnostics, 323107). While this equilibration was occurring, the HybEZ Oven was equilibrated to 40°C. Slides were removed from 5x SSC and washed twice with RNAscope wash buffer. Three drops of the AMP1 were applied to each section and the humidity control tray was placed back in the oven where it incubated for 30 mins at 40°C. Slides were subsequently washed twice with wash buffer and placed back in the slide holder and 3 drops of AMP2 were applied to each section and left to incubate in the oven at 40°C for 30 mins. Slides were washed twice, and treated with 3 drops of AMP3 at 40°C for 15 mins. While the AMP3 incubation was occurring, the dye solutions were prepared as follows. The dyes were prepared at a concentration of 1:1,000 by combining 1000 µl of TSA Buffer (Advanced Cell Diagnostics, Cat#322809) with 1 µL of Opal 520 Reagent (in DMSO, Akoya Biosciences, Cat#OP-001001) and Opal 690 Reagent (in DMSO, Akoya Biosciences, Cat#OP-001006) respectively.

Once the 15-min AMP3 incubation was complete the slides were washed and inserted back into the slide holder, then 3 drops of HRPC1 were applied to each slide, and that was allowed to incubate for 15 mins at 40°C. Once that was complete the slides were washed and returned to the slide holder and 3 drops of the HRP Blocker were added to each section. This was allowed to incubate for 15 mins at 40°C. Slides were washed and 150 µL of the Opal 520 dye mix was added to each section. This was allowed to incubate at 40°C for 30 minutes. This process was then repeated for the C3 channel, which was treated with Opal 690 dye.

Counterstaining Mounting and Imaging: Four drops of RNAscope DAPI (Advanced Cell Diagnostics, Cat#323108) were added to each section for the purpose of counterstaining and left to sit at room temperature for 30 secs. The DAPI was gently tapped off the slide and 50 µL of Prolong Gold antifade reagent (Invitrogen, Cat#P36930) was added inside the barrier but not directly touching the section, avoiding bubbles. A glass coverslip (Corning, 24 x 60 mm, Cat#2975-246) was lowered onto the slide, slides were allowed to dry overnight in a dark slide box (Fisher Brand, Cat#03-448-4) at 4°C before imaging. Slide boxes were stored in the cold room a 4°C for long term storage. Three separate mice were analyzed for each postnatal age, and quantitative data depicts the average values from three mice.

#### Double immunohistochemistry/RNA in situ hybridizations

NeuN or GFAP immunohistochemistry/Adgra1 RNA *in situ* experiments were conducted in the following manner, essentially as described in the manufacturer’s protocol (Advanced Cell Diagnostics Cat#323180 and Cat#323100). Tissue collection, sectioning, and pretreatment were conducted as described above for standard RNA *in situ* experiments up until the initial 10-min room temperature hydrogen peroxide treatment. Following hydrogen peroxide treatment, slides were washed twice with ddH_2_O followed by once with 1x-PBS-T (PBS with 0.1% Tween-20). The slides were returned to the slide holder and 150 µL of the primary antibody (anti-NeuN Mouse, EMD Millipore Corp., Cat#MAB377; anti-GFAP, Invitrogen, Cat#PA1-10004) diluted in RNAscope Co-Detection Antibody Diluent (Advanced Cell Diagnostics, Cat#323160) in a 1:500 concentration was added to each section. Slides were incubated at 4°C overnight in the humidity control tray. Post-primary Fixation and Protease Treatment: After incubation with the primary antibody, slides were washed three times with 1x-PBS-T at room temperature. Then slides were submerged in 10% Neutral Buffered Formalin (Sigma-Aldrich, Cat#65346-85) for 30 mins at room temperature. Following that incubation, slides were washed four times with PBS-T. Slides were subsequently placed back into the humidity control tray, and 4 drops of Protease 4 were added and incubated for exactly 30 mins at room temperature. After incubation the slides were washed three times with ddH_2_O. The RNAscope Multiplex fluorescent assay was then performed, as described above. Following the last HRP blocker step in the RNAscope Multiplex assay, immunofluorescence for NeuN or GFAP was performed. For NeuN, HRP-conjugated goat anti-mouse secondary antibody (Biotium #20400-1mL) was diluted in Co-Detection Antibody Diluent (Advanced Cell Diagnostics, #323160) at a 1:500 concentration was added to completely cover the sections and allowed to incubate at room temperature for 30 mins. For GFAP, goat anti-chicken Alexa Fluor 647 (Invitrogen # A21449) was diluted in Co-Detection Antibody Diluent at a 1:500 concentration and allowed to incubate at room temperature for 30 minutes. Slides were subsequently washed twice with 1x PBS-T. For NeuN, 150 µL of the previously prepared Opal dye (Akoya Biosciences) was added to the slides and incubated for 10 mins at room temperature. Then the slides were washed twice with 1x PBS-T and ready for counterstaining and mounting as described above for standard *in situs*.

#### Confocal Imaging of fixed samples

Images were acquired using a Nikon A1r resonant scanning Eclipse Ti2 HD25 confocal microscope with a 10x (Nikon #MRD00105, CFI60 Plan Apochromat Lambda, N.A. 0.45), 20x (Nikon #MRD00205, CFI60 Plan Apochromat Lambda, N.A. 0.75), and 60x (Nikon #MRD01605, CFI60 Plan Apochromat Lambda, N.A. 1.4) objectives, operated by NIS-24 Elements AR v4.5 acquisition software. Laser intensities and acquisition settings were established for individual channels and applied to entire experiments, and images were collected at the following resolution: 10x -1.73 µm/pixel, 20x - 0.62 µm/pixel, 60x - 0.29 µm/pixel, 60x with deconvolution – 0.07 µm/pixel. Brightness was adjusted uniformly across all pixels for a given experiment for Figure visualization purposes. Images were pseudocolored for Figure visualization purposes. Quantification of fluorescence intensities was conducted by imaging 3-5 image frames per biological replicate, which were averaged to generate a single biological replicate value. The averaged value for each replicate is depicted as open circles in each graph.

#### Image Analysis (Immunocytochemistry)

Images were analyzed using NIS Elements AR 5.42 with a custom-made pipeline in NIS Elements GA3. For synaptic puncta colocalization, the individual channels were processed using a rolling ball radius set to 3.45 µm. Local contrast was enhanced using a radius pool 1.04 and 25% degree. Then, the signal was 3D segmented and thresholded accordingly to detect all visible signal. Next, the signal was subsequently processed with binary option separate (0.035 µm). Lastly, objects were filtered based on size. Exclusion criteria was set 0.010 µm. For each object centroid coordinates for 3 dimensions (x, y, z) were measured in micrometers using the 3D binary processing. Results were exported as a CSV file and further analyzed using MATLAB pdist2 function to calculate the proximity between individual puncta. Colocalization was defined if the proximity between two puncta was less than 500 nm. Percent colocalization was calculated as the proportion of colocalized puncta count relative to total puncta count per image.

#### Image Analysis (Immunohistochemistry)

Images were analyzed using NIS Elements AR 5.42 with a custom-made GA3 pipeline. For synaptic puncta, the individual channels were processed using a rolling ball radius set to 3.45 µm. Local contrast was enhanced with radius pool 1.04 and 25% degree. The signal was then 3D segmented and thresholded accordingly to detect all visible signals. Then the signal was subsequently processed with binary options: smooth and separate. Lastly, objects were filtered based on size with exclusion criteria set to 0.010 µm. With 3D binary processing object count, volume, mean intensity were measured. DAPI 405 channel was processed using a similar pipeline to normalize the measured mean intensity. The 405 channel was processed using Gaussian filter with radius 5, then 3D thresholded accordingly to detect all visible signals. The signals were subsequently processed with binary options: smooth, clean and fill holes. Lastly, objects were filtered based on voxel count with exclusion criteria set to 150. Mean DAPI intensity was calculated by using 3D measurements. Results were exported as a CSV file. For analyzing Synaptotagmin-2 (Syt-2) puncta located on PV soma, the Syt-2 channel was processed using a similar pipeline to determine Syt-2 signal. First, images were converted to maximum intensity projections. Subsequently, the Ai14-546 channel processed with unsharp mask (power 0.50, area 41). Next, intensity was equalized based on histogram of the first frame. The Ai14/546 channel was thresholded to detect soma signal. Then, the segment area was circularly dilated by 0.105 µm and defined as PV cell soma object area. Next, Syt-2 channel was processed using a rolling ball radius set to 3.45 µm. Local contrast was enhanced with radius pool 1.04 and 25% degree. The image was thresholded to detect all visible signal, and a function was used to segment the Syt-2 signal overlapping with the PV cell soma object area. Object count, object intensity and object area were calculated from PV soma Syt-2 signal. The DAPI 405 channel was processed to normalize measured mean object intensity. The 405 channel was processed using a Gaussian filter (radius=1). Then, the signal was thresholded accordingly to detect all visible signal. Lastly, signals were filtered based on area with exclusion criteria set visually. Mean DAPI intensity was calculated, and results were exported as a CSV file.

#### Morphological analysis of granule cells

Biocytin-filled cells were imaged on the Nikon A1r resonant scanning Eclipse Ti2 HD25 confocal microscope as described above. Images for Imaris 3D reconstructions were taken at 60x magnification. X-Y stitches together with Z-stacks were collected to encompass the entire cell at a Z-step size of 0.175 µm. For dendritic spine quantification, each filled cell was imaged at 60x magnification to examine dendritic spines in the molecular layer of the dentate gyrus. Images of multiple dendritic branches in each layer were taken and analyzed to obtain one average spine density value per cell, per region. For 3D Imaris reconstructions, Nikon ND2 files of full cells were converted to Imaris compatible format (.imd) via the Imaris Converter application. Images were then exported to the Imaris Software. Image processing with a default background subtraction level of 206 was used. Filaments were reconstructed by choosing the “autopath no spine function” with only segments within the region of interest (ROI) being assessed. The ROI was defined to ensure the full cell in all 3 dimensions was encompassed. The program was trained through multiple iterations until each seed point was accurately classified. Once classified, segments were trained and predicted through multiple iterations to ensure the entire cell was being reconstructed accurately in all three dimensions. Maximum gap length was set to 1. Reconstructed cells were imaged with a line segment thickness of 5 for ideal viewing. The screenshot feature was used to take the image of the maximum projection of the cell. Quantitative analysis of the cell was exported from the software to Microsoft Excel. Filament Segment Length in microns was the sum of all segment lengths within the entire filament graph. Filament No. of Sholl Intersections was defined as the number of segment intersections on concentric spheres (1.0 µm), defining segment spatial distribution as a function of distance from the beginning point (soma). Filament No. of Segment Branches was calculated by following a segment from the segment beginning point (soma) along all segments for the defined distance, and then counting how many different branches have been reached. Imaris calculates this as: Distances 0 to n * DistanceIncrement, where DistanceIncrement has a default of value of 1 µm and n is dependent on the maximum segment terminal point distance to the beginning point of the filament.

#### Acute slice electrophysiology

Mice were deeply anesthetized with isoflurane, decapitated and their brains were quickly removed and placed into ice cold solution containing (in mM): 228 Sucrose, 2.5 KCl, 1 NaH_2_PO_4_, 26 NaHCO_3_, 0.5 CaCl_2_, 7 MgSO_4_, 11 D-Glucose saturated with 95% O_2_/5% CO_2_. Transverse hippocampal slices (300 μm thick) were cut by a vibratome (Leica VT 1200S) and transferred to a holding chamber containing artificial cerebrospinal fluid (ACSF, in mM): 119 NaCl, 2.5 KCl, 1 NaH_2_PO_4_, 26 NaHCO_3_, 2.5 CaCl_2_, 1.3 MgSO_4_, 11 D-Glucose, ∼290 mOsm. Slices were recovered at 32°C for 30 min, followed by recovery at room temperature for 1 hr in the same holding chamber. Acute slices were transferred to a recording chamber continuously perfused with oxygenated ACSF (1.5 ml/min) maintained at 32°C. Whole cell recordings were performed from granule cell layer of the dentate gyrus. For whole-cell patch-clamp experiments, the patch pipettes were pulled from borosilicate glass capillary tubes (World Precision Instruments, Cat#TW150-4) using a PC-100 pipette puller (Narishige PC-100). The resistance of pipettes filled with whole cell pipette solution varied between 3-5 MΩ. Synaptic currents were monitored with a Multiclamp 700B amplifier (Molecular Devices) synchronized with Clampex 11.2 data acquisition software (Molecular Devices). Electrophysiological data were digitized with Digidata 1550B (Molecular Devices). The recording rig contained a Nikon Eclipse FN1 microscope controlled via NIS Elements software with 4X (CFI60 Plan Fluor 4X objective lens, N.A. 0.13) and 40X (CFI60 Apochromat 40X near infrared water dipping lens, N.A. 0.8), pco Edge 4.2 LT sCMOS camera (Cat#77067009), and Sutter micromanipulators (MPC-200). For voltage-clamp recordings of excitatory transmission, a whole-cell pipette solution was used containing (in mM) 135 Cs-Methanesulfonate, 8 CsCl, 10 HEPES, 0.25 EGTA, 0.3 Na2GTP, 2 MgATP, 7 phosphocreatine, 0.1 Spermine (pH 7.3, adjusted with CsOH and 302 mOsm). For voltage-clamp recordings of inhibitory transmission, a whole cell pipette solution was used containing (in mM) 146 CsCl, 10 HEPES, 0.25 EGTA, 2 MgATP, 0.3 Na2GTP, 7 phosphocreatine, 0.1 Spermine (pH 7.3, adjusted with CsOH and 296 mOsm). The external bath solution contained (in mM) 140 NaCl, 5 KCl, 2 CaCl2, 0.8 MgCl2, 10 HEPES, and 10 glucose (pH 7.35, adjusted with NaOH). AMPAR- and NMDAR-excitatory postsynaptic currents (EPSCs) were pharmacologically isolated by adding the γ-aminobutyric acid receptor blocker picrotoxin (50 µM; Tocris, Cat#1128) to the extracellular bath solution. Inhibitory postsynaptic currents (IPSCs) were isolated by adding AP5 (50 µM; Tocris Cat#0106) and CNQX (10 µM; Tocris, Cat#1045) to the extracellular bath solution to block NMDA and AMPA receptors, respectively. Spontaneous miniature postsynaptic currents (mEPSCs and mIPSCs) were monitored at room temperature in the presence of tetrodotoxin (1 µM; Tocris, Cat#1069) to block action potential triggered neurotransmitter release. Whole cell patch clamp experiments were done in patch configuration while holding the cells at -70 mV. Synaptic currents were sampled at 10 kHz and analyzed offline using Clampfit 11.2 software (Molecular Devices). Miniature events were monitored for 5 min 10 s and the last 4 min 45 s of each trace was analyzed. Miniature events were analyzed using the template matching search and a minimal threshold of 5 pA, and each event was visually inspected for inclusion or rejection. For voltage-clamp recordings of evoked postsynaptic currents (eIPSCs) QX-314 (1 mM; Tocris, Cat#1014), a blocker of voltage gated Na+ channels, was added to the previously described inhibitory whole cell patch pipette solutions. Local stimulation was provided for eIPSCs recordings using a concentric bipolar electrode (FHC, Cat#CBAEB75) immersed into external bath solution and controlled by Sutter micromanipulator (MPC-200). The concentric bipolar electrode was placed in the granule cell layer and was kept at a consistent distance (150 µm) with minimal variability from each cell to deliver stimulation to the field. Stimulation intensities were set to 50-100-200-400 µA in increasing order. For measuring paired-pulse ratio (PPR), paired pulses with 400 µA intensity were delivered with the bipolar electrode with the following order of intervals: 25-50-100-250-500 ms. Five sweeps were measured for each inter-stimulus interval. Between interval changes, 30 sec period was allowed for the recovery of the neurotransmitter pool. For biocytin fill, internal solution was made as above except with the addition of 2 mg/mL Biocytin (Sigma, Cat#B4261). After the recordings, the patch pipette was gently removed, and the slices were transferred to PBS in a 24 well plate. The PBS was immediately exchanged with 4%PFA/PBS and slices were incubated overnight at 4°C, followed by 5 x 5 min washes with PBS. Samples were permeabilized for 30 min in 0.3% Triton X-100/PBS at room temperature, blocked for 1 hr at room temperature in 5% normal goat serum/0.1% Triton X-100/PBS, and incubated with Streptavidin Alexa Fluor™ 647 conjugate (ThermoFisher, Cat#S21374) diluted 1:1,000 into blocking buffer for 90 min at room temperature. Samples were labeled with DAPI (Sigma, Cat#10236276001) diluted into PBS for 5 mins. Samples were subsequently washed 4 x 5 min with PBS and mounted as described for immunohistochemistry.

#### Mouse behavior

Previously mentioned ADGRA1 PV and SST cKO mouse lines were used for the behavior assays. All mice were aged to 12 weeks, and approximately equal numbers of male and female mice were used. Control and cKO mice were randomly distributed across cages to ensure randomization during testing. Experimenters were blinded to the genotype of the mice before, during and after the testing. All animals were housed in a temperature- and humidity-controlled housing facility and were kept on a 12:12 hr light cycle. All procedures were approved by the Vanderbilt Institutional Animal Care and Use Committee. All behavioral assays were conducted at the Vanderbilt Murine Neurobehavioral Laboratory Core. Mice were acclimated to the facility at least two weeks before the experiments. Running order of the assays was kept consistent and run at the same time each day between 7 a.m. and noon. Assays were conducted in the following order: Open field assay, fear conditioning, seizure induction.

#### Open Field Assay

Exploratory locomotor activity was measured in specially designed chambers measuring 27 x 27 cm (ENV-510; MED Associates, Georgia, VT, USA). Chambers were housed in sound-attenuating cases to restrict auditory stimuli of the surroundings during testing. Infrared beams and detectors were used to record the movement in the open field arena automatically. Locomotor activity was measured over 60 mins. Overall activity in the box, rearing count, time and distance traveled in the center area of the box were measured. Total rearing count was measured automatically by beam breaks. Time spent exploring center area (19.05 × 19.05 cm) versus periphery (50% of surface area) of the chamber was used as a proxy for measuring anxiety profile.

#### Fear conditioning assay

Mice were placed in a sound-attenuating chamber with a wire grid floor capable of transmitting an electric shock. Their movements were monitored by cameras fixed to the doors. On the first day, mice were trained to associate an auditory tone with a small electric shock. During training trials (total duration 8 mins) a 30 sec tone was played at the end of which a small shock was administered (2 sec, 0.5 mA). The tone shock pairing was repeated during the training trial 3 times. On the subsequent test day, the mouse was exposed to the same chamber for a 4-min trial (no tone, no shock). In this context-retrieval trial time spent immobile was reported as time freezing. One hr later, a different context was created to conduct the cue retrieval (memory of the tone) trial. Room lighting was changed to ambient red light. The chamber was altered using a white curved plexiglass wall and floor insertion, and a 10% vanilla smell was placed in an inaccessible container. Following a 2 min exploration of the novel context the tone was played for 2 mins, and the freezing response was measured. Freezing behavior was monitored automatically by software (VideoFreeze, Med Associates, USA) and time spent immobile was reported as time freezing.

#### PTZ-induced seizure severity assay

Pentylenetetrazol (PTZ) (Sigma-Aldrich, Cat#P6500-25G) was freshly reconstituted on the day of the assay in normal saline to a concentration of 5 mg/mL. Mice were weighed immediately before the assay, and the appropriate dose was calculated. PTZ was administered via intraperitoneal (i.p.) injection. Two doses were selected for PTZ: 35 mg/kg and 50 mg/kg, representing the low and high doses. Doses were selected based on the ranges commonly reported in comparable studies. Behavioral observations and video recordings were started immediately after PTZ administration and continued for 30 mins. After the test period, mice were either returned to their home cages or euthanized. Cages were not returned to the housing facility until at least 3 hrs of monitoring after drug administration, following confirmation that no further ictal phases were observed. If status epilepticus was observed (continuous seizure activity longer than 90 secs) assay was ended with isoflurane and mice were euthanized. Seizure severity was scored according to a modified Racine scale (Table 2). Behavioral recordings were divided into 5-min intervals over the 30-min observation period. Each interval was scored based on the highest seizure activity that observed. Behavioral scoring was performed by an experimenter who was blinded to the experimental conditions of the mice. For seizure latency measurements, the time of onset of the first observable seizure behavior was recorded.

**Table 2.** Modified Racine Scale.

| Severity Score | Behavioral Description |
| --- | --- |
| 0 | Normal behavior, no abnormality. |
| 1 | Immobilization, lying on belly. |
| 2 | Head nodding, facial, forelimb, or hindlimb myoclonus. |
| 3 | Continuous whole-body myoclonus, myoclonic jerks, tail held up stiffly. |
| 4 | Rearing, tonic seizure, falling on its side. |
| 5 | Tonic-clonic seizure, falling on its back, wild rushing and jumping |
| 6 | Seizure resulting with sudden death |

## Acknowledgements

We thank Jennifer Skelton and Leesa Sampson at the Vanderbilt Genome Editing Facility for generating Adgra1 floxed mice. We thank Fiona Harrison and Krista Paffenroth for expertise in mouse behavioral studies. We thank Oleg Kovtun at the Cell Imaging Shared Resource core for expertise with image analysis. We thank all members of the Sando laboratory for critical feedback on the study. This study was supported by grants from the NIH (R00-MH117235 and DP2MH1401324 to RS) and Alfred Sloan Foundation (Sloan Fellowship in Neuroscience to RS).

## Author Contributions

B. Tosun performed all electrophysiological, immunohistochemical, and RiboTag experiments. B. Tosun conducted behavioral analysis in conjunction with the Vanderbilt Mouse Behavioral Facility. E. Orput conducted RNA *in situ* experiments. D. Bui performed BRET2 studies. R. Sando conducted molecular cloning. B. Tosun, E. Orput, D. Bui, and R. Sando performed analysis and interpretation. B. Tosun and R. Sando wrote the manuscript.

## SUPPLEMENTAL DATA FIGURES and FIGURE LEGENDS

**Figure S1:**
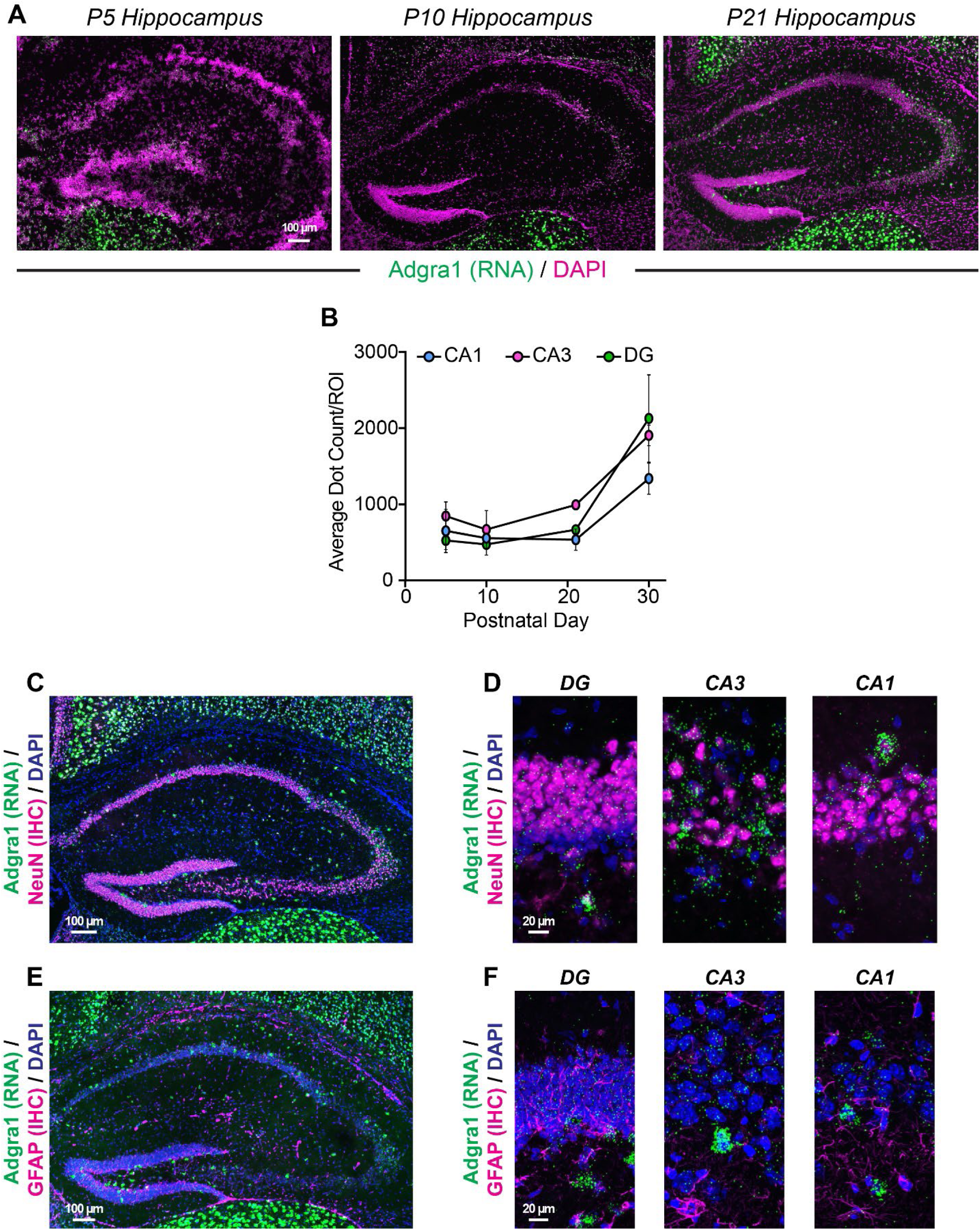
Developmental timecourse of *Adgra1* expression in the mouse hippocampus. **A,** representative RNA *in situ* images for *Adgra1* at postnatal day 5 (P5), P10, or P21. **B,** quantification of average *Adgra1* RNA puncta per region-of-interest (ROI) in the DG, CA1, or CA3 at the indicated developmental time points. **C-F,** analysis of *Adgra1*-expressing cells with double RNA *in situ* hybridization/immunohistochemistry. **C,** representative hippocampal section labeled for *Adgra1* RNA *in situ* probe together with immunohistochemistry for the neuronal marker NeuN. **D,** high-magnification images from experiments in **C** in the DG, CA3, or CA1. **E & F,** analysis of *Adgra1* expression in glia. Representative RNA *in situs* for *Adgra1* which were co-labeled via immunohistochemistry for GFAP. Numerical data are means ± SEM. See Figure 1 for studies at P30.

**Figure S2:**
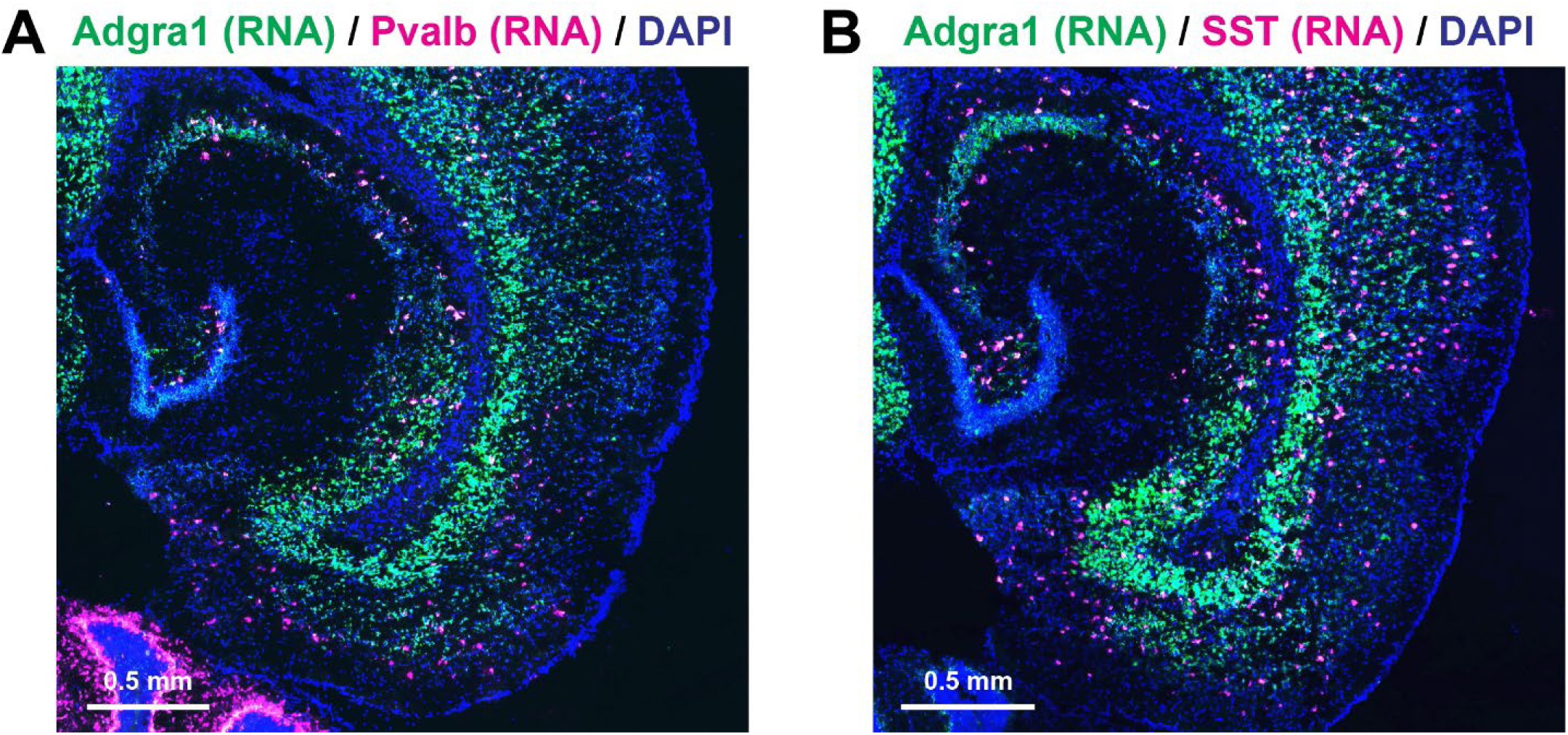
Horizontal sections depicting *Adgra1* expression in the entorhinal cortical-hippocampal circuit at postnatal day 30 (P30) **A,** *Adgra1/Pvalb* RNA *in situs* in horizontal sections from P30 mice. **B,** similar to **A**, except for *Adgra1/SST*. See Figure 2 for additional studies of *Adgra1* expression in hippocampal interneurons.

**Figure S3:**
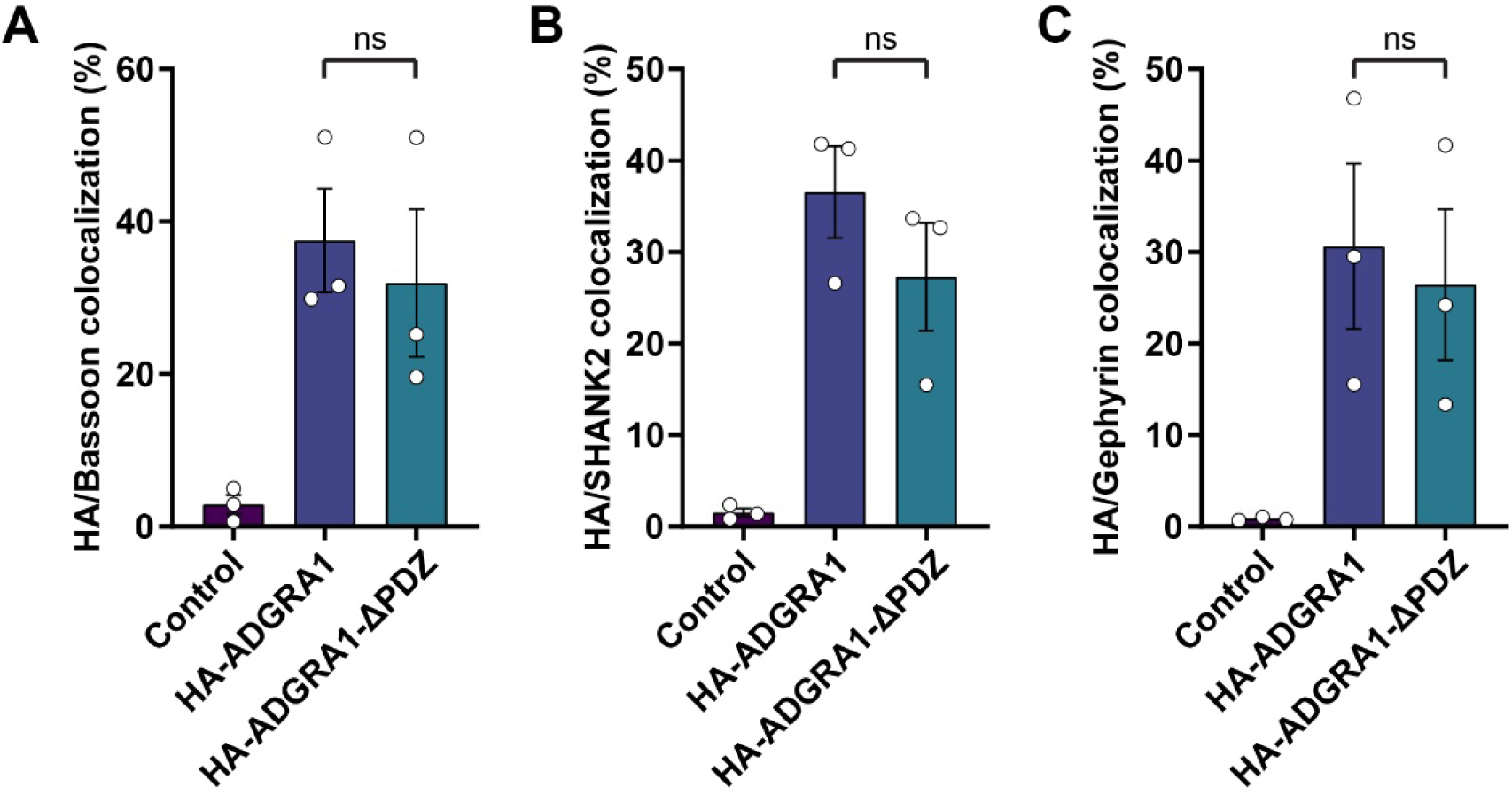
Quantification of HA-ADGRA1 colocalization with synaptic markers. **A,** Co-localization of lentiviral HA-ADGRA1 or a PDZ truncation (HA-ADGRA1-ΔPDZ) and presynaptic Bassoon. **B,** similar to **A**, except for excitatory postsynaptic SHANK2. **C,** similar to **A**, except for inhibitory postsynaptic Gephyrin. Numerical data are means ± SEM. Statistical significance was assessed via one-way ANOVA with *post hoc* Tukey tests. See Methods for details on co-localization analysis.

**Figure S4:**
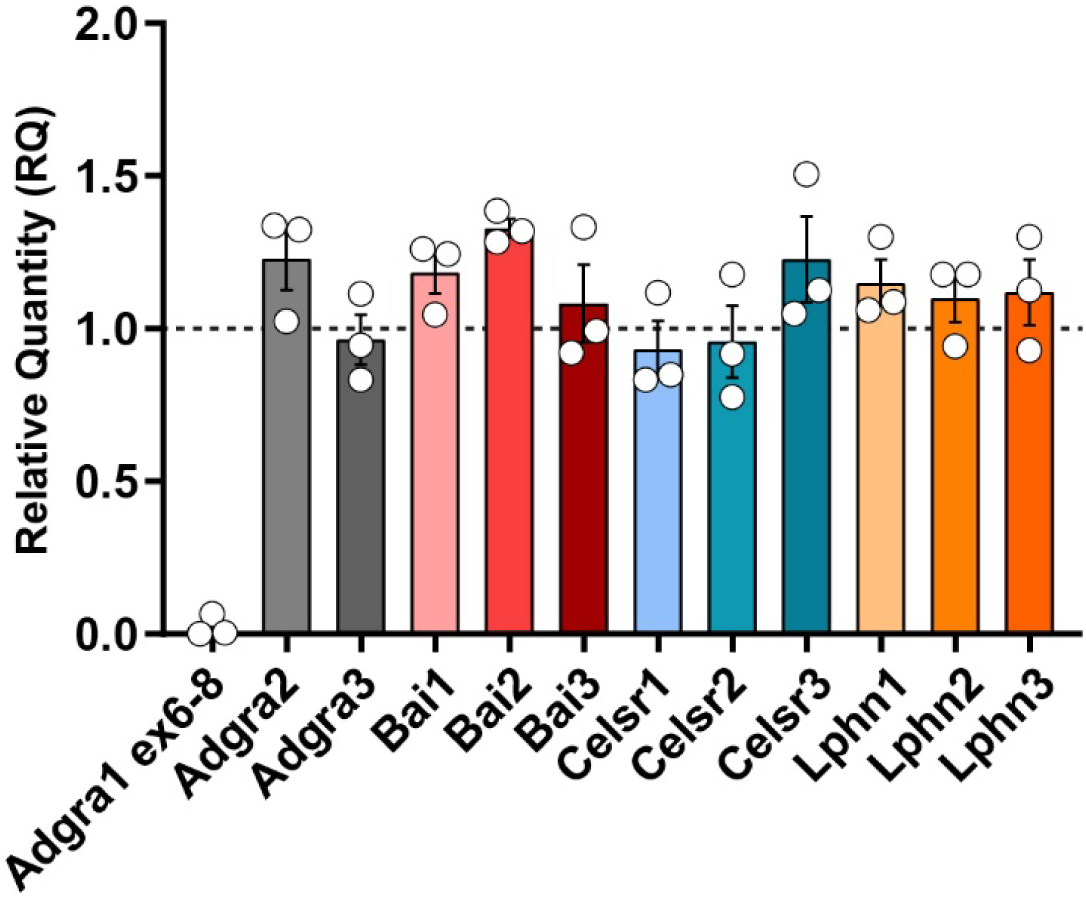
Validation of Adgra1 cKO mice. RT-qPCR analysis with indicated qPCR probes. Primary hippocampal cultures were generated from Adgra1 homozygous floxed mice and infected with lentiviruses encoding either CRE or inactive ΔCRE. Adgra1 cKO mice harbor floxed exon 6. Data was normalized to *Gapdh* levels. Numerical data are means ± SEM from 3 independent culture replicates.

**Figure S5:**
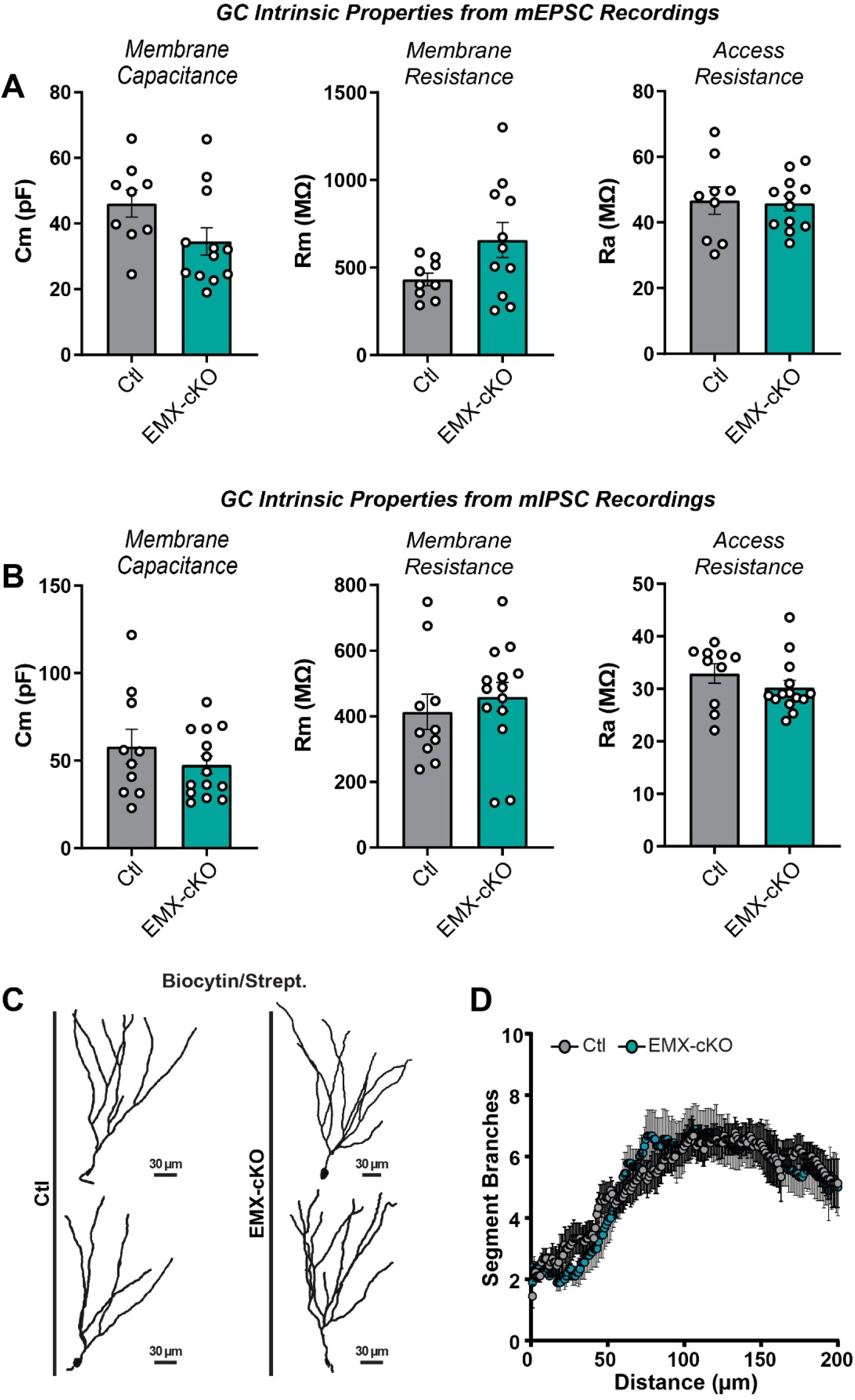
Additional DG GC properties from studies in Figure 3. **A,** membrane capacitance, membrane resistance, and access resistance measurements from mEPSC GC recordings in Figure 3E-G. **B,** membrane capacitance, membrane resistance, and access resistance measurements from mIPSC GC recordings in Figure 3H-J. **C & D,** analysis of dendritic arborization in DG GCs. **C,** representative 3D reconstructions of DG GCs following biocytin loading and streptavidin labeling. **D,** average segment branches at indicated distances from DG GC soma. Numerical data are means ± SEM. See Figure 3 for analysis of mEPSC/mIPSC frequency and amplitude and analysis of dendritic spine density from biocytin-filled DG GCs.

**Figure S6:**
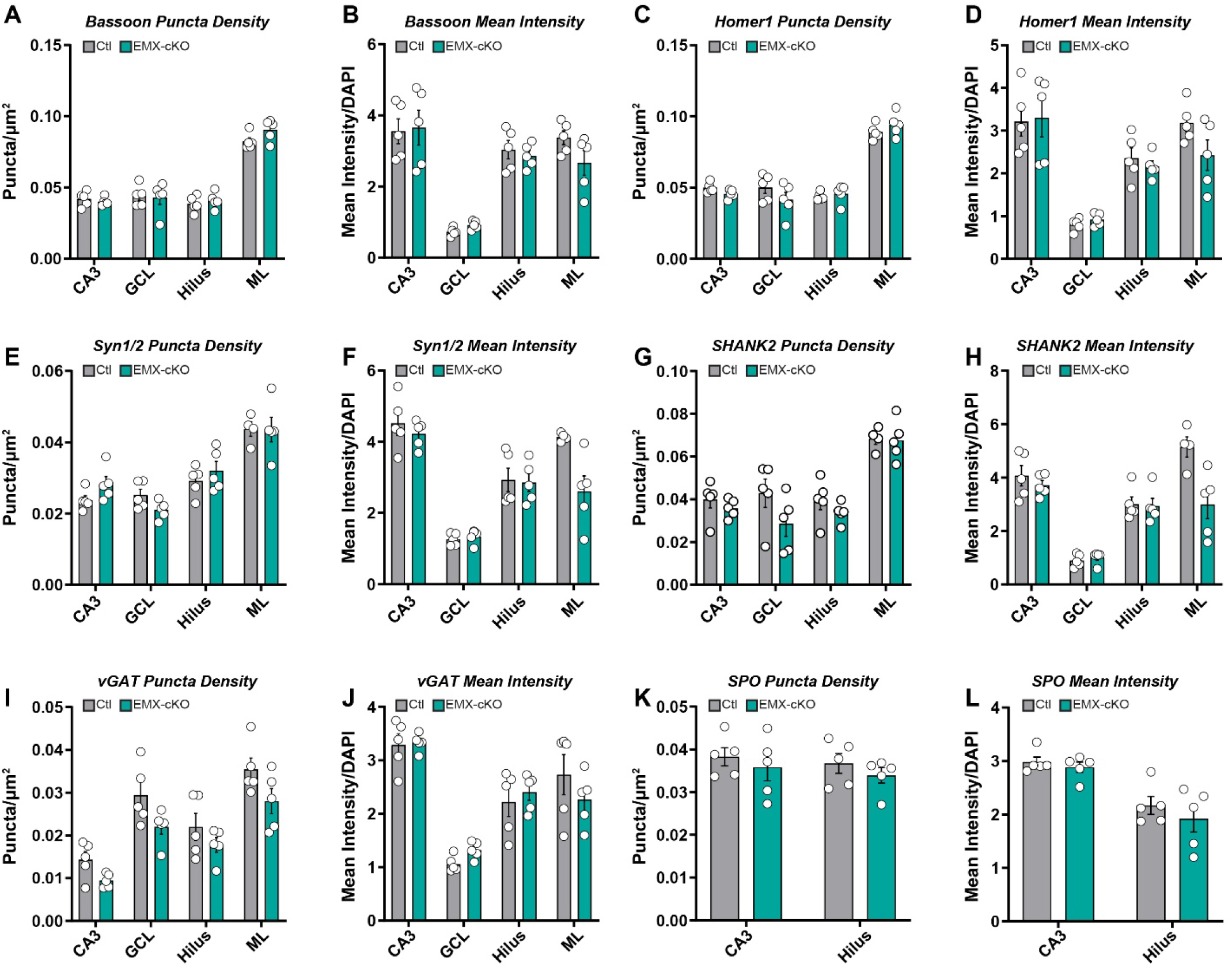
Quantification of synapse density and mean fluorescence intensity from experiments in Figure 3K-M. **A & B,** quantification of Bassoon puncta density (**A**) and mean intensity relative to DAPI intensity as an internal control (**B**) in indicated hippocampal sub-regions. **C & D,** similar to **A** and **B**, except for Homer1. **E & F,** similar to **A** and **B**, except for Syn1/2. **G & H,** similar to **A** and **B**, except for SHANK2. **I & J,** similar to **A** and **B**, except for vGAT. **K & L,** analysis of Synaptoporin (SPO), a marker of large mossy fiber terminals (LMTs), puncta density (**K**) or mean intensity relative to DAPI (**L**) in the CA3 or hilus. Numerical data are means ± SEM from 5 mice per genotype. Statistical significance was assessed via 2-way ANOVA. See Figure 3K-M for representative images.

**Figure S7:**
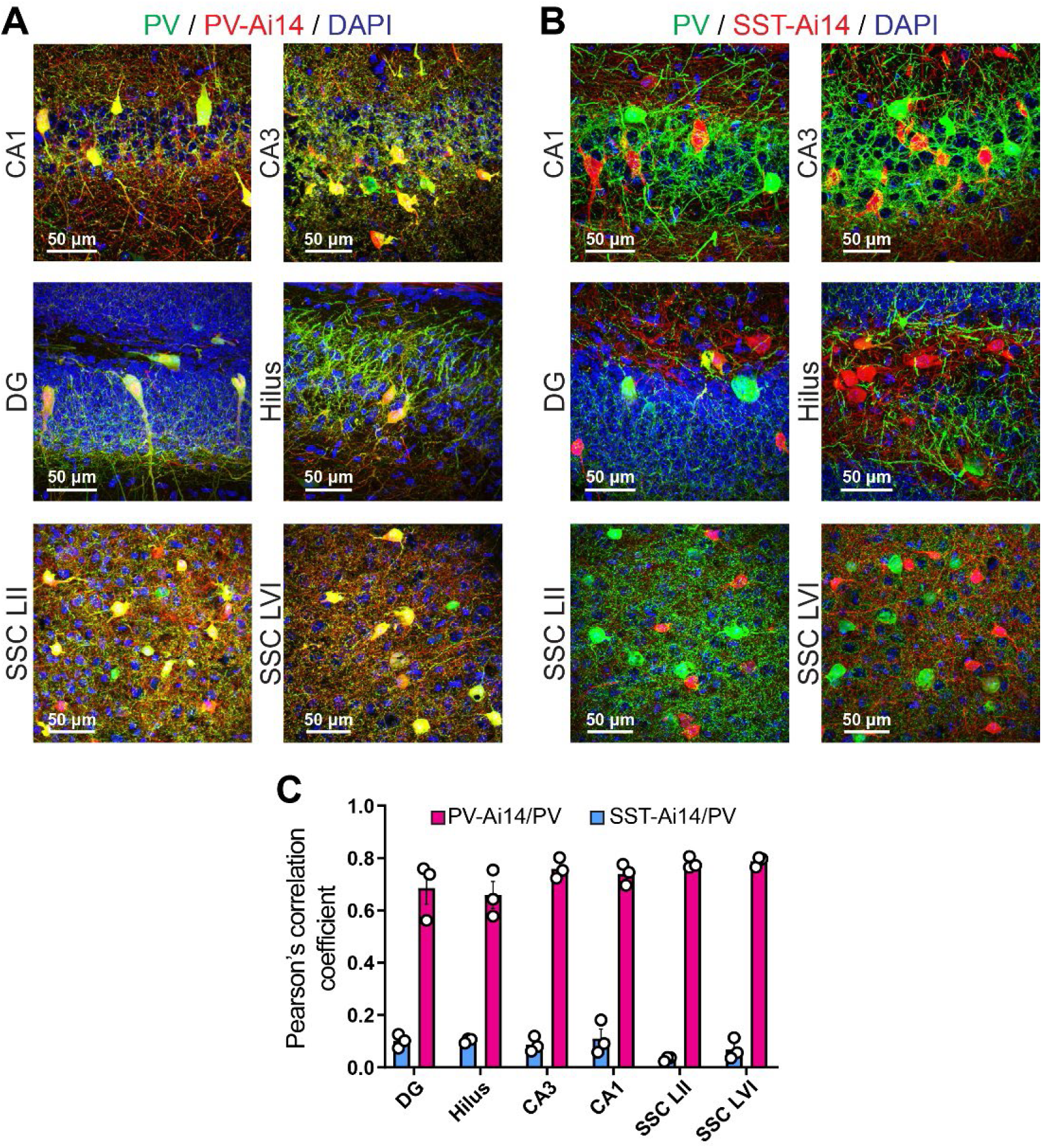
Analysis of the specificity of PV-Cre and SST-Cre driver lines. **A,** representative images of sections from PV-Cre/Ai14 mice immunolabeled for endogenous PV. **B,** similar to **A**, except for sections from SST-Cre/Ai14 mice. **C,** quantification of Pearson’s correlation coefficient between the Ai14 and PV immunostaining channels in the indicated brain regions. Numerical data are means ± SEM from 3 mice per genotype.

**Figure S8:**
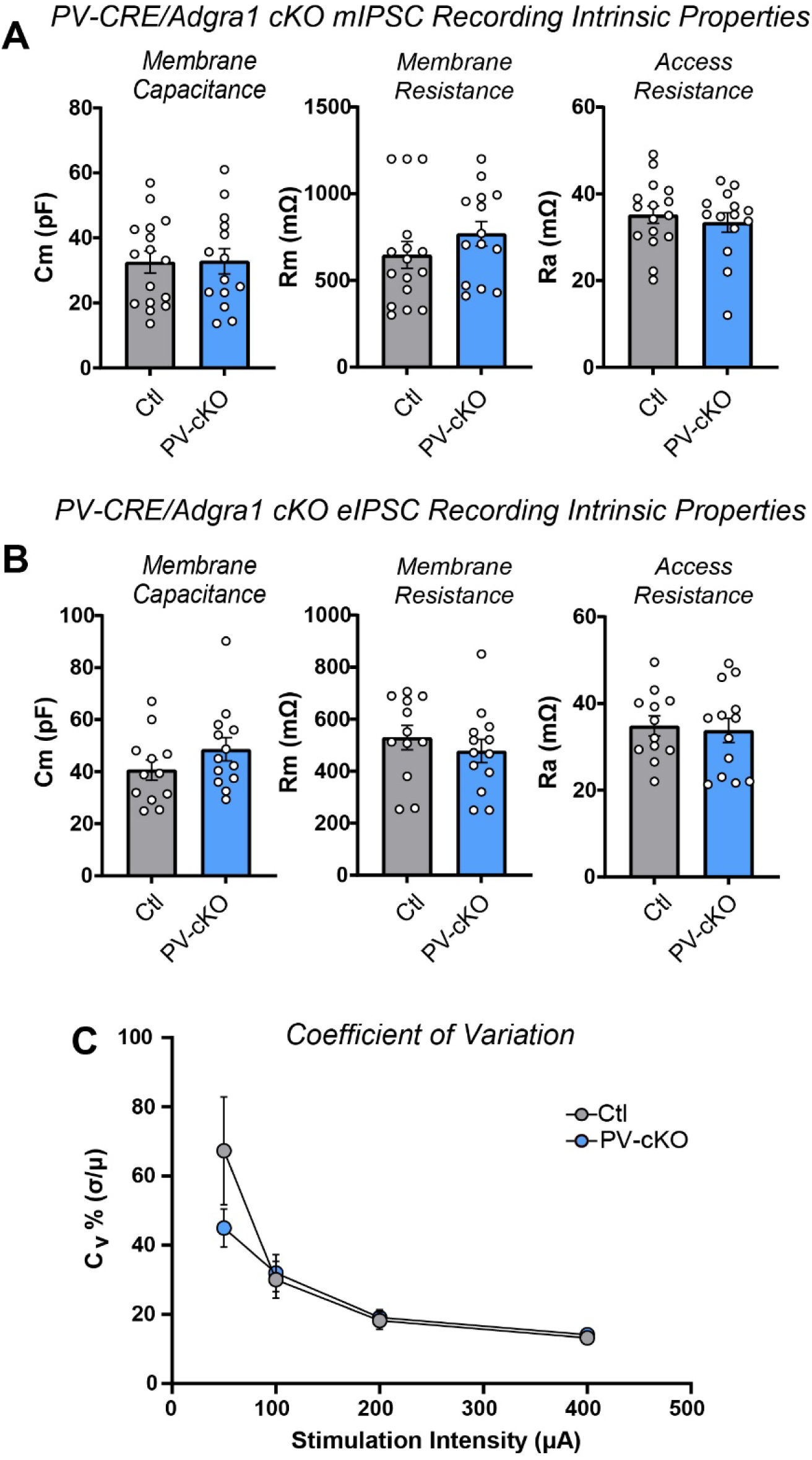
Intrinsic properties of GCs from recordings in Figure 4. **A,** membrane capacitance, membrane resistance, and access resistance measurements from mIPSC GC recordings in Figure 4B-D. **B,** membrane capacitance, membrane resistance, and access resistance measurements from eIPSC GC recordings in Figure 4E-H. **C,** analysis of coefficient of variation from eIPSC recordings in Figure 4E & F. Numerical data are means ± SEM. See Figure 4 for analysis of mIPSC frequency and amplitude and eIPSC amplitude and paired-pulse ratios.

**Figure S9:**
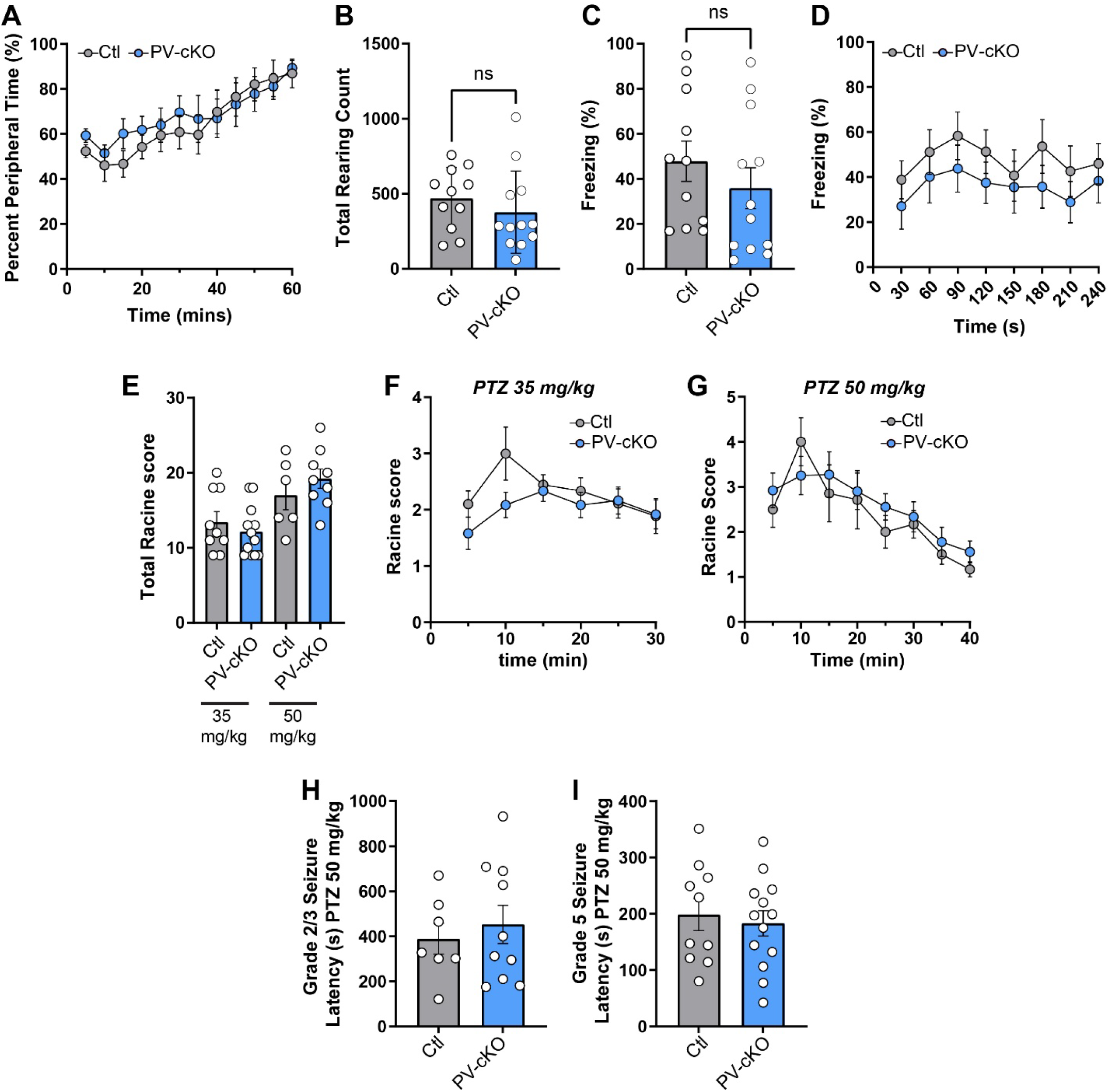
Additional behavioral characterization of PV-cKO lines. **A,** percent time spent in the peripheral 50% over 60-minute open field trials. **B,** quantification of average total rearing counts during the open field assay. **C & D,** contextual fear conditioning studies. **C,** percent time spent freezing in the context conditioned stimulus following fear conditioning. **D,** percent time spent freezing over time within the conditioned context chamber. **E-I,** analysis of PTZ-induced seizure susceptibility. **E,** cumulative total Racine score following either a moderate (35 mg/kg) or high (50 mg/kg) dose of PTZ. **F,** Racine score over time following an intraperitoneal injection of 35 mg/kg PTZ. **G,** similar to **F**, except for 50 mg/kg PTZ. **H & I,** latency to grade 2/3 (**H**) or grade 5 (**I**) seizure onset following 50 mg/kg PTZ injections. Numerical data are means ± SEM with individual mice depicted as open circles. See Figure 5 for additional behavioral analyses. Statistical significance was determined via two-tailed t-test, one-way ANOVA with *post hoc* Tukey test, or two-way ANOVA.

**Figure S10:**
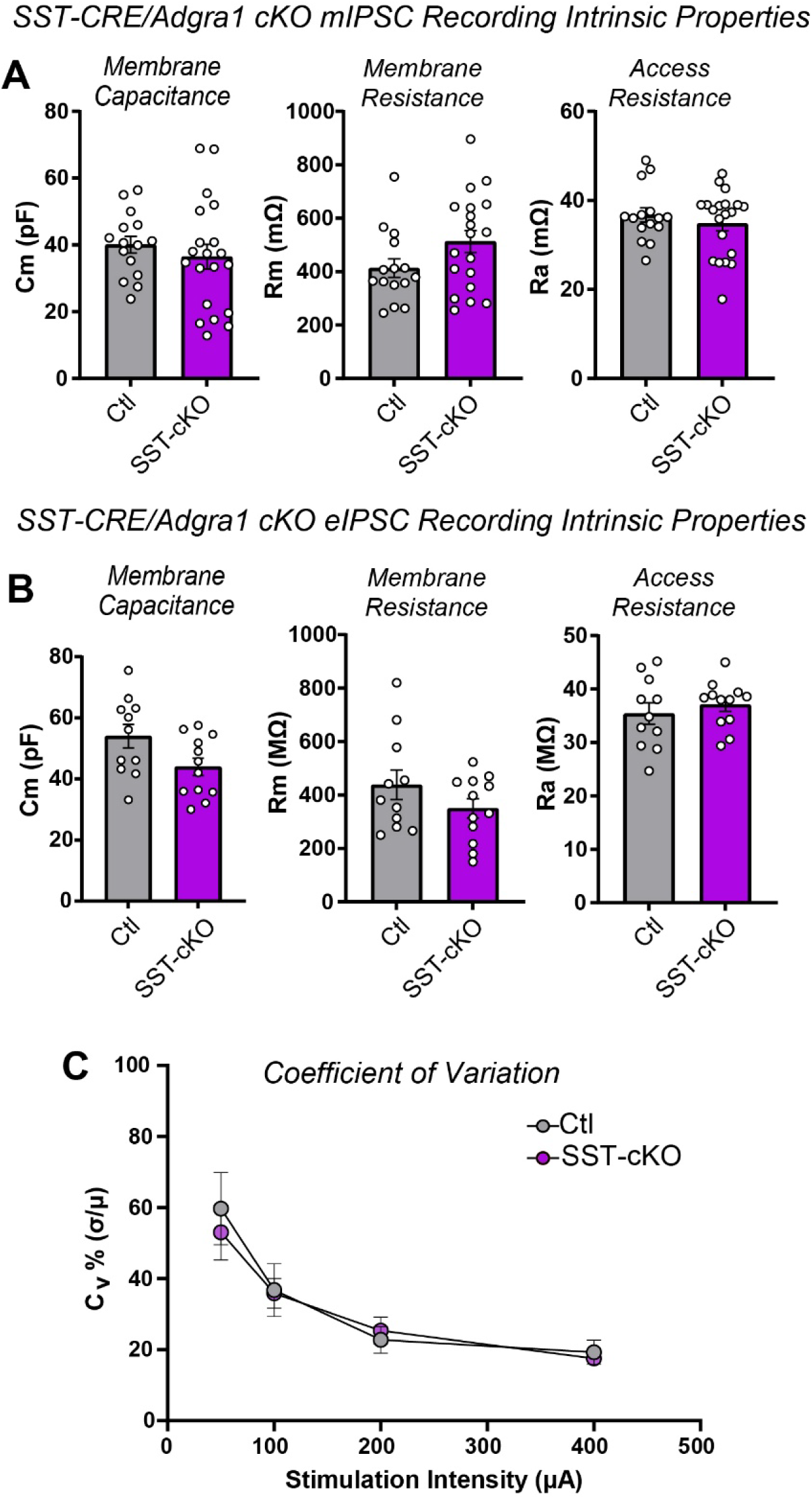
Additional electrophysiological properties from Figure 6. **A,** membrane capacitance, membrane resistance, and access resistance measurements from mIPSC GC recordings in Figure 6B-D. **B,** membrane capacitance, membrane resistance, and access resistance measurements from eIPSC GC recordings in Figure 6E-H. **C,** analysis of coefficient of variation from eIPSC recordings in Figure 6E & F. Numerical data are means ± SEM. See Figure 6 for analysis of mIPSC frequency and amplitude.

**Figure S11:**
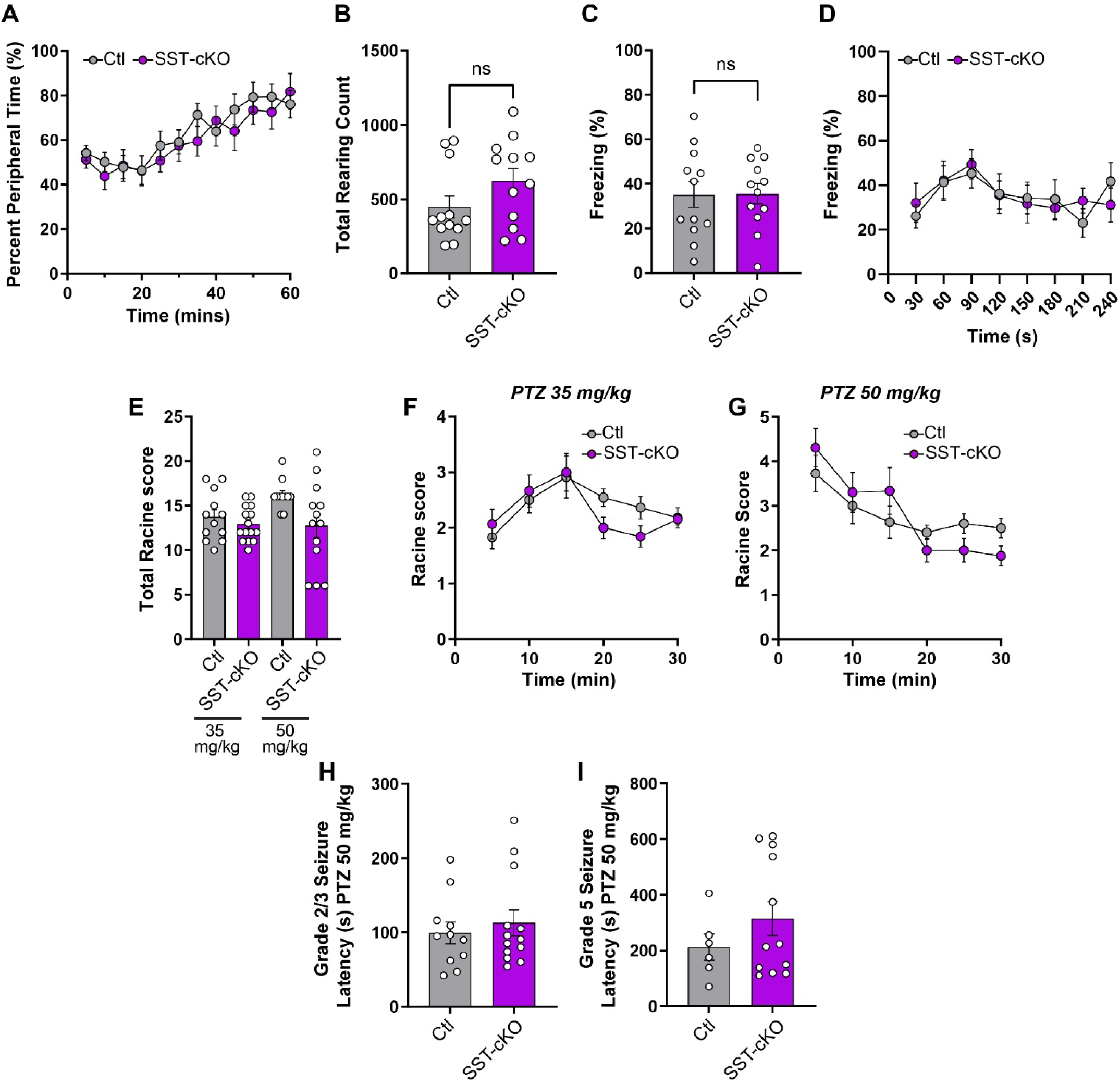
Additional behavioral analysis from Figure 6. **A,** percent time spent in the peripheral 50% over 60-minute open field trials. **B,** quantification of average total rearing counts during the open field assay. **C & D,** contextual fear conditioning studies. **C,** percent time spent freezing in the context conditioned stimulus following fear conditioning. **D,** percent time spent freezing over time within the conditioned context chamber. **E-I,** analysis of PTZ-induced seizure susceptibility. **E,** cumulative total Racine score following either a moderate (35 mg/kg) or high (50 mg/kg) dose of PTZ. **F,** Racine score over time following an intraperitoneal injection of 35 mg/kg PTZ. **G,** similar to **H**, except for 50 mg/kg PTZ. **H & I,** latency to grade 2/3 (**H**) or grade 5 (**I**) seizure onset following 50 mg/kg PTZ injections. Numerical data are means ± SEM. Statistical significance was determined via two-tailed t-test, one-way ANOVA with *post hoc* Tukey test, or two-way ANOVA.

**Figure S12:**
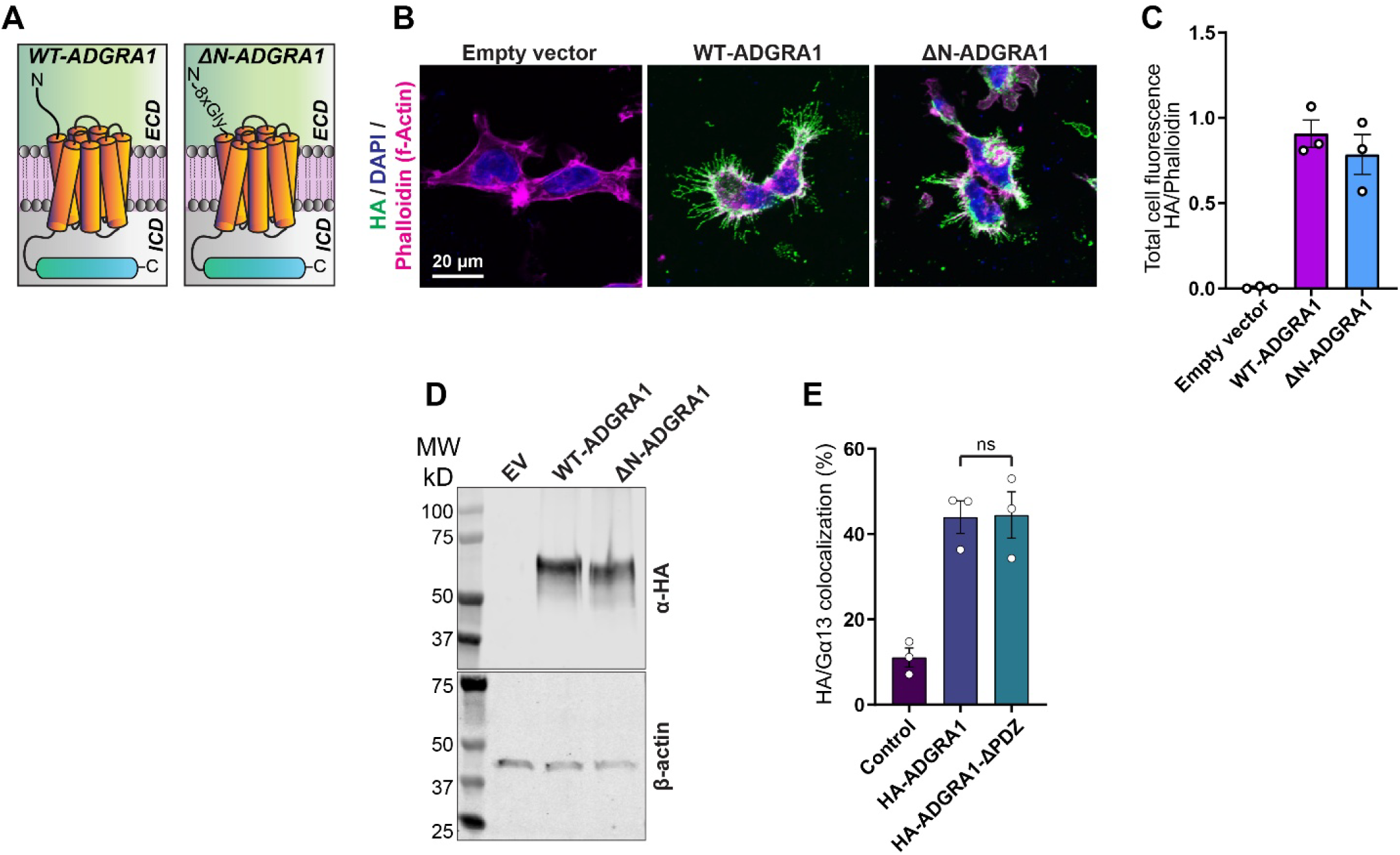
Analysis of ADGRA1 constructs from Figure 7. **A,** diagram of WT ADGRA1 and the ΔN-ADGRA1 mutant, which contained an 8x Glycine linker substituting the short ADGRA1 extracellular sequence. **B,** representative images of HEK293T cells transfected with indicated experimental conditions and labeled for HA tag together with Phalloidin (f-Actin) and DAPI as controls. **C,** quantification of HA intensity relative to Phalloidin intensity, which was used as an internal control. **D,** representative immunoblot for HA tag or β-actin in indicated experimental conditions. **E,** co-localization of lentiviral HA-ADGRA1 or a PDZ truncation (HA-ADGRA1-ΔPDZ) and Gα13. Numerical data are means ± SEM from 3 independent replicates. See Figure 7 for TRUPATH BRET2 analysis of ADGRA1 G protein activation.

